# High Frequencies of Antiviral Effector Memory T_EM_ Cells and Memory B Cells Mobilized into Herpes Infected Vaginal Mucosa Associated With Protection Against Genital Herpes

**DOI:** 10.1101/2023.05.23.542021

**Authors:** Nisha Rajeswari Dhanushkodi, Swayam Prakash, Afshana Quadiri, Latifa Zayou, Mahmoud Singer, Nakayama Takashi, Hawa Vahed, Lbachir BenMohamed

**Affiliations:** Laboratory of Cellular and Molecular Immunology, Gavin Herbert Eye Institute, University of California Irvine, School of Medicine, Irvine, CA 92697; Department of Molecular Biology and Biochemistry; Institute for Immunology; the University of California Irvine, School of Medicine, Irvine, CA 92697; Kindai University, Higashiosaka, Osaka 577-8502, Japan; Department of Vaccines and Immunotherapies, TechImmune, LLC, University Lab Partners, Irvine, CA 92660; USA

**Keywords:** HSV-2, genital herpes, CCL28, mucosa, memory CD8^+^ T cells, memory B cells

## Abstract

Vaginal mucosa-resident anti-viral effector memory B- and T cells appeared to play a crucial role in protection against genital herpes. However, how to mobilize such protective immune cells into the vaginal tissue close to infected epithelial cells remains to be determined. In the present study, we investigate whether and how, CCL28, a major mucosal-associated chemokine, mobilizes effector memory B- and T cells in leading to protecting mucosal surfaces from herpes infection and disease. The CCL28 is a chemoattractant for the CCR10 receptor-expressing immune cells and is produced homeostatically in the human vaginal mucosa (VM). We found the presence of significant frequencies of HSV-specific memory CCR10^+^CD44^+^CD8^+^ T cells, expressing high levels of CCR10 receptor, in herpes-infected asymptomatic (ASYMP) women compared to symptomatic (SYMP) women. A significant amount of the CCL28 chemokine (a ligand of CCR10), was detected in the VM of herpes-infected ASYMP B6 mice, associated with the mobilization of high frequencies of HSV-specific effector memory CCR10^+^CD44^+^ CD62L^-^ CD8^+^ T_EM_ cells and memory CCR10^+^B220^+^CD27^+^ B cells in the VM of HSV-infected asymptomatic mice. In contrast, compared to wild-type (WT) B6 mice, the CCL28 knockout (CCL28^(-/-)^) mice: (*i*) Appeared more susceptible to intravaginal infection and re-infection with HSV-2; (*ii*) Exhibited a significant decrease in the frequencies of HSV-specific effector memory CCR10^+^CD44^+^ CD62L^-^ CD8^+^ T_EM_ cells and of memory CD27^+^B220^+^ B cells in the infected VM. The results imply a critical role of the CCL28/CCR10 chemokine axis in the mobilization of anti-viral memory B and T cells within the VM to protect against genital herpes infection and disease.

## INTRODUCTION

Genital herpes caused by herpes simplex virus type 1 and type 2 (HSV-1 and HSV-2) affects over 490 million (13%) people 15–49 years of age worldwide (1). Over the past several decades, considerable efforts have been made to develop a herpes simplex vaccine, but such a vaccine remains an unmet medical need (2). This results in a significant global health and financial burden. Approximately forty to sixty million individuals are infected with HSV-2 in the United States alone, with nearly six to eight hundred thousand reported annual clinical cases (3–8). HSV-2 and HSV-1 replicate predominantly in the mucosal epithelial cells and establish latency in the sensory neurons of the dorsal root ganglia (DRG) where, in symptomatic individuals, they reactivate sporadically causing recurrent genital herpetic disease (4, 9, 10). Both HSV-1 and HSV-2 cause genital herpes disease through infection of the mucosa of the genital tract. Genital herpes can produce genital ulcers increasing the risk of acquiring and transmitting HIV infection (11–13).

In response to HSV-1 and HSV-2 infections, the vaginal epithelial cells secrete soluble factors including chemokines that mobilize and guide leukocytes of the innate and adaptive immune system, such as the NK cells, neutrophils, monocytes, B and T cells to the site of infection, vaginal mucosa (VM), or DRG the site of reactivation. Apart from their role in the mobilization of immune cells, chemokines can signal through specific membrane-bound receptors that lead to the activation of cellular pathways that can eliminate the virus. Out of all 48 known human chemokines, CCL25, CCL28, CXCL14, and CXCL17 mucosal chemokines are especially important in mucosal immunity because they are homeostatically expressed in mucosal tissues (14–17). The chemokine expression in the vagina mucosa influences the mobilization and activation of innate immune cells that facilitate adaptive immune responses (4, 9).

Local B and T cell responses within the VM play an important role in the defense against herpes infection and disease (18–22)*. However, the VM tissue appears to be immunologically restricted and mostly resistant to accepting homing B and T cells that could be traveling from the draining lymph nodes and circulation.* (23–26). Major gaps within the current literature include the identity of involved chemokines and the underlying mechanisms through which these chemokines and their receptors mobilize the protective memory B and T cell subsets into the infected and inflamed vaginal mucosal tissues. Several chemokines are produced in the vaginal mucosa following genital HSV-2 infection (23–26), but whether and how these chemokines affect mucosal B and T cell responses in the vaginal mucosa remains to be fully elucidated.

In this study, we first performed bulk RNA sequencing of HSV-specific CD8^+^ T cells to determine any differential regulation of the chemokine pathways in HSV-infected symptomatic (SYMP) asymptomatic vs. (ASYMP) women. Subsequently, we identified the CCL28, also known as mucosae-associated epithelial chemokine (MEC), (a chemoattractant for CCR10 expressing B and T cells), as being highly expressed in HSV-infected ASYMP women. Moreover, using the CCL28 knockout mouse model, we confirmed the role of the CCL28/CCR10 chemokine axis in protective B and T cell immunity against genital herpes. In this report, we demonstrate the role of the CCL28/CCR10 chemokine axis in the mobilization of circulating B and T memory cells into the VM site of infection and the underlying CCL28/CCR10 chemokine axis-mediated mechanism of action. In this study, we discussed the potential use of the mucosal chemokine CCL28 to improve genital herpes immunity and protect against infection and disease caused by HSV, and potentially other sexually transmitted viruses.

## MATERIALS AND METHODS

### Virus propagation and titration

Rabbit skin (RS) cells (from ATCC, VA, USA) grown in Minimum Essential Medium Eagle with Earl’s salts and L-Glutamine (Corning, Manassas, VA) supplemented with 10% fetal bovine serum and 1% penicillin-streptomycin was used for virus propagation. HSV-2 strain186 was propagated in RS cells as described previously (20–22). The virus was quantified by plaque assay in RS cells. The HSV-2 strain 186 was originally isolated from a genital lesion from an individual attending a sexually transmitted disease clinic in Houston, Texas, in the 1960s. Strain 186 is used in this study as it is a highly pathogenic herpes virus (87).

### Mice

Female C57BL/6 (B6) wild-type mice (6-8 weeks old) were purchased from the Jackson Laboratory (Bar Harbor, ME) and CCL28^(-/-)^ KO mice breeders were a kind donation by Dr. Takashi Nakayama, Kindai University, Japan). CCL28^(-/-)^ KO mice breeding was conducted in the animal facility at UCI where female mice at 6-8 weeks were used. Animal studies conformed to the Guide for the Care and Use of Laboratory Animals published by the US National Institute of Health. Animal studies were conducted with the approval of the Institutional Care and Use Committee of the University of California-Irvine (Irvine, CA) and conformed to the Guide for the Care and Use of Laboratory Animals published by the US National Institute of Health (IACUC protocol #19-111).

### Genital infection of mice with HSV-2

All animals were injected subcutaneously with 2mg progesterone (Depo-Provera^®^), to synchronize the ovarian cycle and increase susceptibility to herpes infection, and then received an IVAG HSV-1 challenge. Previous studies have shown that estrogen might have a crucial role in the protection against genital infection by regulating MEC/CCL28 expression in the uterus (58). Since immune responses in the VM compartment appear to be under the influence of sex hormones, future studies will compare the phase of the menstrual cycle/estrous cycle in mice as well as in symptomatic and asymptomatic women. Mice were intravaginally infected with 5 x 10^3^ pfu of HSV-2 strain 186 in 20μL sterile PBS. Following genital infection, mice were monitored daily for genital herpes infection and disease progression. For genital inflammation and ulceration examination, pictures were taken at the time points listed in the figure legends using a Nikon D7200 camera with an AF-S Micro NIKKOR 105mm f/2.8 lens and a Wireless Remote Speedlight SB-R200 installed. CCL28 KO and WT that survived the primary infection were re-infected with 5 x 10^3^ pfu HSV-2 strain186 at day 30 p.i. At day 10 post-re-infection, mice were euthanized and immune cells from VM and spleen were used for flow cytometry. Single-cell suspensions from the mouse vaginal mucosa (VM) after collagenase treatment (15mg/ml) for 1 hour were used for FACS staining.

### Monitoring of genital herpes infection and disease scoring in mice

Virus shedding was quantified in vaginal swabs collected on days 3, 5, 7, and 10 p.i. Infected mice were swabbed using moist type 1 calcium alginate swabs and frozen at -80 LJC until titrated on RS cell monolayers, as described previously (30–34). Mice were scored every day from day 1 to day 9 p.i for pathological symptoms. Stromal keratitis was scored as 0-no disease; 1-cloudiness, some iris detail visible; 2-iris detail obscured; 3-cornea opaque; and 4-cornea perforation. Mice were evaluated daily and scored for epithelial disease (erythema, edema, genital ulcers, and hair loss around the perineum) and neurological disease (urinary and fecal retention and hind-limb paresis/paralysis) on a scale that ranged from 0 (no disease) to 4 (severe ulceration, hair loss, or hind-limb paralysis) (88, 89). Mice that reached a clinical score of 4 were euthanized.

### Bulk RNA sequencing on sorted CD8^+^ T cells

RNA was isolated from the sorted CD8^+^ T cells using the Direct-zol RNA MiniPrep (Zymo Research, Irvine, CA) according to the manufacturer’s instructions. RNA concentration and integrity were determined using the Agilent 2100 Bioanalyzer. Sequencing libraries were constructed using TruSeq Stranded Total RNA Sample Preparation Kit (Illumina, San Diego, CA). Briefly, rRNA was first depleted using the RiboGone rRNA removal kit (Clonetech Laboratories, Mountain View, CA) before the RNA was fragmented, converted to double-stranded cDNA and ligated to adapters, amplified by PCR, and selected by size exclusion. Following quality control for size, quality, and concentrations, libraries were multiplexed and sequenced to single-end 100-bp sequencing using the Illumina HiSeq 4000 platform.

### Differential gene expression analysis

Differentially expressed genes (DEGs) were analyzed by using integrated Differential Expression and Pathway analysis tools. Integrated Differential Expression and Pathway analysis seamlessly connect 63 R/Bioconductor packages, two web services, and comprehensive annotation and pathway databases for homo sapiens and other species. The expression matrix of DEGs was filtered and converted to Ensemble gene identifiers, and the preprocessed data were used for exploratory data analysis, including *k*-means clustering and hierarchical clustering. The pairwise comparison of symptomatic and asymptomatic groups was performed using the DESeq2 package with a threshold of false discovery rate < 0.5. and fold change >1.5. Moreover, a hierarchical clustering tree and network of enriched GO/KEGG terms were constructed to visualize the potential relationship. Gene Set Enrichment Analysis (GSEA) method was performed to investigate the related signal pathways activated among symptomatic and asymptomatic groups. The Parametric Gene Set Enrichment Analysis (PSGEA) method was applied based on data curated in Gene Ontology and KEGG. The pathway significance cutoff with a false discovery date (FDR) ≥ 0.2 was applied.

### Flow cytometry

Single-cell suspensions from the mouse VM after Collagenase D (Millipore Sigma, St. Louis, MO) treatment (15mg/ml) for 1h at 37C were used for FACS staining. The following antibodies were used: anti-mouse CD3 (clone 17A-2, BD Biosciences), CD45 (clone 30-F11, BD Biosciences), CD4, CD8, CD44, CD62L, B220and CD27 (BD Biosciences). For surface staining, mAbs were added against various cell markers to a total of 1 x 10^6^ cells in phosphate-buffered saline containing 1% FBS and 0.1% Sodium azide (fluorescence-activated cell sorter [FACS] buffer) and left for 45 minutes at 4°C. Cells were washed again with FACS buffer and fixed in PBS containing 2% paraformaldehyde (Sigma-Aldrich, St. Louis, MO).

### HSV-2-specific ASC ELISPOT assay

Immune cells isolated from VM of HSV-2 infected mice (2 million cells/ ml) were stimulated in B-cell media containing mouse polyclonal B cell activator (Immunospot) for 5 days. CTL Mouse B-Poly-S are stock solutions containing Resiquimod and either recombinant Human IL-2 or recombinant Mouse IL-2 respectively, used for the polyclonal expansion of memory B cells Subsequently, cells were washed in RPMI medium and plated in specified cell numbers in ELISPOT membrane plates coated with heat-inactivated HSV-2. The ASC-secreting cells were detected after 48 hours of the addition of cells to ELISPOT plates. The ELISpot plates were detected by imaging using an ELISPOT reader (ImmunoSpot). The spots were detected and quantified manually.

### Immunohistochemistry for human VM tissue

For immunohistochemistry, human vaginal mucosa sections were used for CCL28 staining. Sections were deparaffinized and rehydrated before the addition of primary antibody anti-human CCL28 for overnight incubation. HRP-labeled secondary antibodies (Jackson Immunoresearch, PA) were used before the addition of substrate DAB. Hematoxylin was used for counterstaining these slides. Subsequently, after thoroughly washing in PBS 3 times slides were mounted with a few drops of mounting solution. Images were captured on the BZ-X710 All-in-One fluorescence microscope (Keyence).

### Virus titration in vaginal swabs

Vaginal swabs (tears) were analyzed for viral titers by plaque assay. RS cells were grown to 70% confluence for plaque assays in 24-well plates. Transfer medium in which vaginal swabs were stored was added after appropriate dilution at 250 ul per well in 24-well plates. Infected monolayers were incubated at 37°C for 1 hour, rocked every 15 minutes for viral adsorption, and then overlaid with a medium containing carboxymethyl cellulose. After 48 hours of incubation at 37°C, cells were fixed and stained with crystal violet, and viral plaques were counted under a light microscope. Positive controls were run with every assay using our previously tittered laboratory stocks of McCrae.

### Statistical analysis

Data for each assay were compared by ANOVA and Student’s *t*-test using GraphPad Prism version 5 (La Jolla, CA). As we previously described, differences between the groups were identified by ANOVA and multiple comparison procedures (33, 34). Data are expressed as the mean ± SD. Results were considered statistically significant at a *P* value of ≤ 0.05.

## RESULTS

### 1. Increased expression of CCR10, the receptor of CCL28 chemokine, on HSV-specific CD8^+^ T cells from herpes-infected asymptomatic women compared to symptomatic women

We first determined whether there are differential expressions of chemokine and chemokine receptor pathways in HSV-specific CD8^+^ T cells from herpes-infected symptomatic women compared to symptomatic women. CD8^+^ T cells specific to HSV-2 gB_561-569_ and VP11-12_220-228_ epitopes were sorted from PBMC of HSV-infected SYMP and ASYMP women and subjected to bulk-mRNA sequencing. As shown in **Fig. 1A** major chemokine and chemokine receptor-specific pathways were significantly upregulated among HSV-infected ASYMP women compared to HSV-infected SYMP women (*P* < 0.05) (**Supplementary Table 1**). In **Figs. 1B** and **1C**, particularly, both the heatmaps (*top panels*) and the volcano plots (bottom *panels*) showed a significant upregulation of CCR10, the receptor of CCL28 chemokine, in CD8^+^ T cell-specific to HSV-2 gB_561-569_ epitope (**Fig. 1B**) and HSV-2 VP11-12_220-228_ epitope (**Fig. 1C**) isolated from ASYMP women, compared to SYMP women. Using flow cytometry, we confirmed high frequencies of CCR10 expressing immune cells in HSV-infected ASYMP women (*n* = 9) compared to HSV-1 infected SYMP women (*n* = 9) (**Fig. 1D**). There was a significant increase in frequencies of CCR10 positive lymphocytes detected in HSV-infected ASYMP women compared to low frequencies of CCR10 positive lymphocytes in SYMP women (i.e., 4.9% vs. 2.6%, *P* = 0.04, Fig. 1D top panels). Moreover, higher frequencies of CCR10^+^CD8^+^ T cells, but not of CCR10^+^CD4^+^ T cells, were detected in HSV-2 infected ASYMP women as compared to HSV-2 infected SYMP women (0.61% vs. 0.27%, *P* = 0.03, Fig. 1D bottom panels). High levels of CCL28 chemokine expression were found in the epithelial cells of the VM in HSV-2-infected women. As detected by immunohistochemistry, the CCL28 is specifically expressed within the Stratum Corneum (SC) and Sub layer of the epithelium in the human VM (**Fig. S1**).

**Figure 1.**
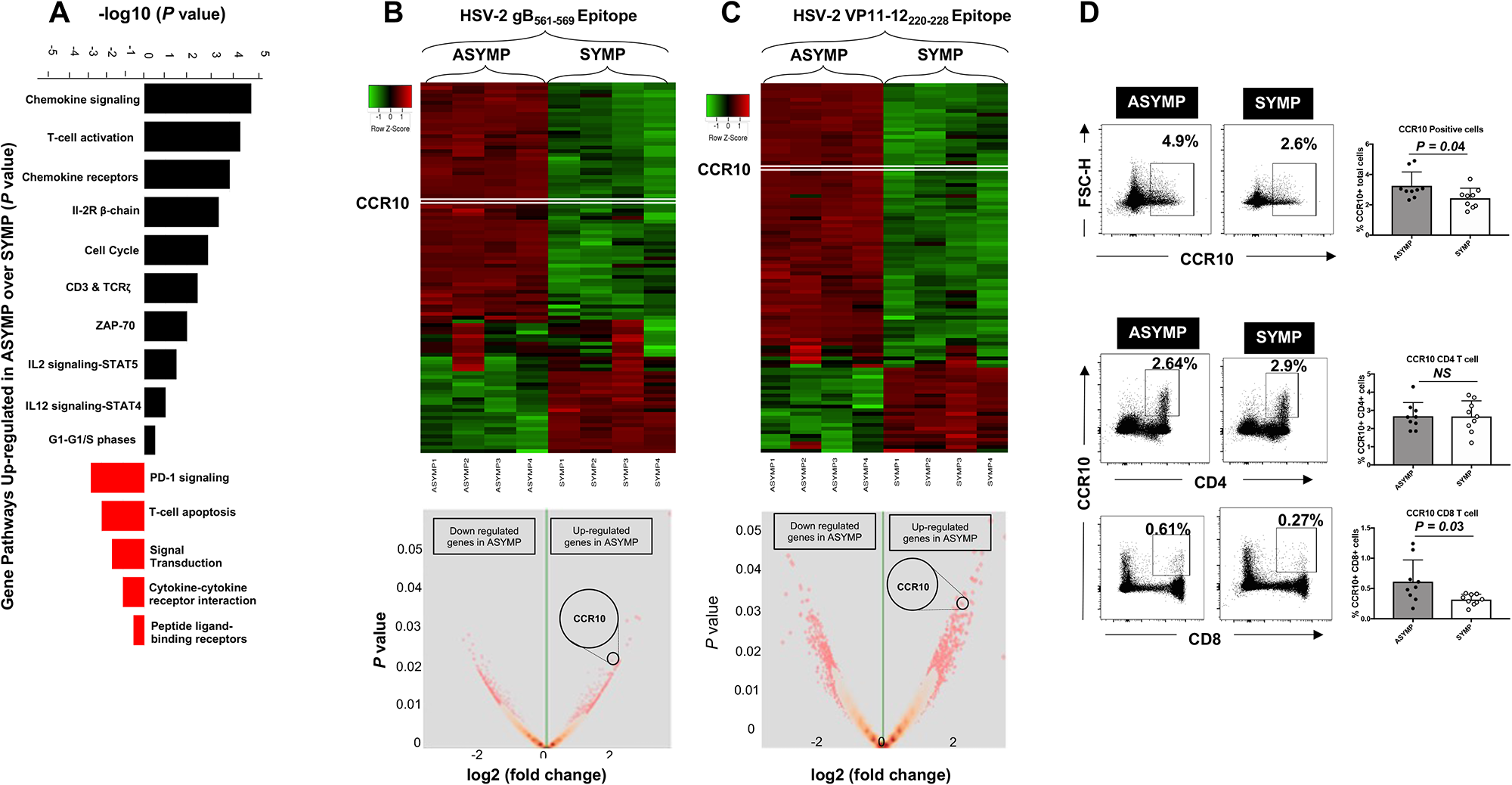
CCR10 expression level in CD8^+^ T cells from PBMC of herpes-infected SYMP compared to ASYMP patients. (**A**) Major gene-specific pathways detected in CD8^+^ T cells from PBMC of herpes-infected SYMP compared to ASYMP patients. (**B**) Differential gene expression (DGE) analysis using bulk RNA sequencing for HSV-2 gB_561-569_ epitope-specific CD8^+^ T cells from SYMP (*n* = 4) vs. ASYMP patients (*n* = 4) shown as a heatmap (*top panel*) and a volcano plot (*bottom panel*). (**C**) Differential gene expression (DGE) analysis using bulk RNA sequencing for VP11-12_220-228_ epitope-specific CD8^+^ T cells from SYMP vs. ASYMP patients heatmap (*top panel*) and a volcano plot (*bottom panel*). (**D**) Representative dot plots showing the frequency of CCR10 in total lymphocytes from SYMP compared to ASYMP patients (*top left panel*). Average frequencies of CCR10 in total lymphocytes from PBMC of SYMP (*n* = 9) and ASYMP (*n* = 9) HSV-1 infected patients (*top right panel*). Representative dot plots showing the frequency of CCR10^+^CD4^+^ T cells and CCR10^+^CD8^+^ T cells from SYMP compared to ASYMP patients (*bottom left panels*). Average frequencies of CCR10^+^CD4^+^ T cells and CCR10^+^CD8^+^ T cells from PBMCs of SYMP (*n* = 9) and ASYMP (*n* = 9) HSV-1 infected patients (*bottom right panel*s). The results are representative of two independent experiments. The indicated *P* values are calculated using the unpaired t-test, comparing results obtained from SYMP vs. ASYMP patients.

Altogether, these results indicate a significant upregulation of CCR10, the receptor of CCL28 chemokine, on HSV-specific CD8^+^ T cells is associated with asymptomatic genital herpes. Additionally, **Supplementary Table 1** shows the differential gene expression (DGE) in HSV-specific CD8^+^ T cells from herpes-infected symptomatic women compared to symptomatic women.

### 2. The CCL28 chemokine is highly produced in the vaginal mucosa of HSV-2-infected B6 mice and is associated with asymptomatic genital herpes

We next determined whether the CCL28 chemokine would be associated with the protection against genital herpes seen in HSV-infected asymptomatic (ASYMP) mice following genital infection with HSV-2. B6 mice (*n* = 20) were infected intra-vaginally (IVAG) with 2 x 10^5^ pfu of HSV-2 (strain MS) (**Fig. 2A**). The vaginal mucosa (VM) was harvested at day 14 post-infection (dpi) and cell suspensions were assayed by flow cytometry for the frequencies of CD8^+^ T cells expressing CCR10, the receptor of CCL28 among total cells (**Fig. 2B**). The level of CCL28 was compared in the VM cell extracts from (*i*) HSV-infected symptomatic (SYMP) mice; (*ii*) HSV-infected asymptomatic (ASYMP) mice; and (*iii*) non-infected control (naive) mice, using ELISA, Immunohistochemical (IHC), and western blot (**Fig. 2C** to **2E**). As shown in **Fig. 2B**, there was a significant increase in the frequency of CCR10^+^CD8^+^ T cells expressing CCR10, the receptor of CCL28 in the VM of ASYMP HSV-infected B6 mice (*HSV-2*) compared to the SYMP HSV-infected B6 mice (*P* = 0.002). Moreover, increased levels of CCL28 chemokine were detected by ELISA quantification in the VM extracts of HSV-2-infected ASYMP mice as compared to HSV-2-infected SYMP mice (**Fig. 2C**). We confirmed an increased expression of CCL28 in HSV-2-infected ASYMP mice compared to HSV-2-infected SYMP mice by the IHC staining of VM sections (**Fig. 2D**) and by Western blot analysis of VM lysates (**Fig. 2E**).

**Figure 2.**
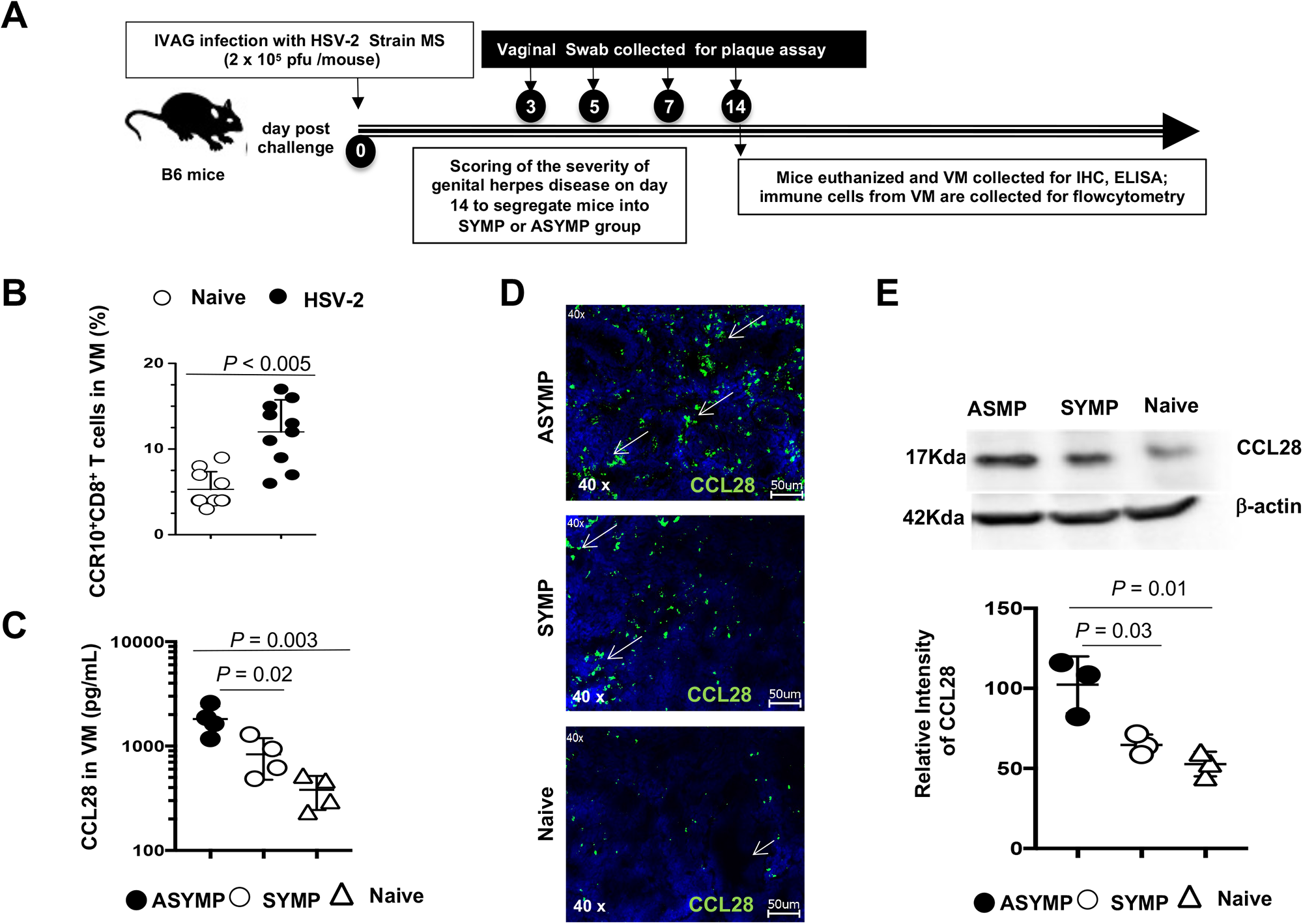
Production of CCL28 chemokines in the vaginal mucosa of HSV-2-infected SYMP and ASYMP B6 mice. (**A**) Experimental plan showing B6 mice (*n* = 20) were infected intra-vaginally (IVAG) with 2 x 10^5^ pfu of HSV-2 (strain MS). The severity of genital herpes disease was scored for 14 days to segregate mice into SYMP or ASYMP groups, as described in the *Material and Methods.* On day 14 post-infection (dpi), SYMP and ASYMP mice and non-infected naïve mice (controls were euthanized and the vaginal mucosae were harvested and cell extracts were assayed by flow cytometry for frequencies of CD8^+^ T cells expressing CCR10, the receptor of CCL28 (i.e., CCR10^+^CD8^+^ T cells), and for CCL28 chemokine using IHC and ELISA. (**B**) Frequency of CCR10^+^CD8^+^ cells among total VM cells determined by flow cytometry in individual HSV-infected ASYMP (n = 4), SYMP (*n* = 4), and control non-infected (*naïve*) (*n* = 8) B6 mice. (**C**) The level of CCL28 chemokine quantified by ELISA (Abcam kit: *ab210578*) in the VM lysates of HSV-infected symptomatic (*SYMP*) B6 mice, HSV-infected asymptomatic B6 mice (*ASYMP*), and non-infected control B6 mice (*Naïve*). VM lysates from each mouse (*n* = 3) were pooled for this experiment. (**D**) Immunohistochemical staining of CCL28 (green) and DAPI (blue) in VM sections harvested on day 8 post-infection (dpi), from ASYMP, SYMP, and Naïve B6 mice. The lower panel shows a graph summarizing the fluorescence intensity (quantitated using Fiji) for CCL28 in the VM of mice. (**E**) Immunoblot of VM lysates from ASYMP, SYMP, and Naïve B8 mice (*n* = 3) probed using western blot for CCL28 (Abcam mAb clone ab23155) (*top panel*). The relative intensity of CCL28 normalized to b-actin is shown in the *bottom panel*. The results are representative of two independent experiments. The indicated *P* values are calculated using the unpaired t-test, comparing results obtained in SYMP vs. ASYMP and results obtained in ASYMP vs. Naïve mice.

Altogether, these results indicate that: (*i*) The intravaginal infection with HSV-2 mobilized higher frequencies of CCR10^+^CD8^+^ T cells expressing CCR10, the receptor of CCL28 in the VM of infected B6 mice; and (*ii*) A significant production of the CCL28 chemokine in the VM of HSV-2 infected B6 mice is associated with asymptomatic genital herpes. These results suggest a role for the CCL28/CCR10 chemokine axis in the protection against symptomatic genital herpes.

### 3. CCL28 deficiency is associated with severe genital herpes and increased virus replication following intravaginal HSV-2 re-infection

To further substantiate the role of the CCL28 chemokine in genital herpes immunity, we studied the functional consequences of CCL28 deficiency in protection against genital herpes infection and disease in mice. CCL28 knockout mice (CCL28*^(-/-)^* mice) and WT mice (*n* =12) were IVAG infected on day 0 with 5 x 10^3^ pfu of HSV-2 (strain 186) (**Fig. 3A**). Mice were scored every day for 14 days p. I for signs of genital herpes and the severity of genital herpes scored, as described in *Material and Methods* (**Fig. 3A**). The disease was scored as 0-no disease, 2-swelling and redness of external vagina, 3-severe swelling and redness of vagina and surrounding tissue and hair loss in the genital area, 4-ulceration and hair loss in the genital and surrounding tissue. Vaginal swabs were collected on days 3, 5, and 7 p.i. to determine virus titers (**Fig. 3A**). As shown in **Fig. 3B**, following primary HSV-2 infection, there was no significant difference detected in the severity of genital herpes between CCL28*^(-/-)^* and WT mice at day 8 p.i. and no significant difference observed in the survival of CCL28*^(-/-)^* and WT mice following IVAG infection with HSV-2 (**Fig. 3C**). In addition, we did not detect any significant difference in virus replication detected in the vaginal swabs collected at day 2, 5, and 7 post-infection from CCL28^-/-^ and WT mice following IVAG infection with HSV-2 (**Fig. 3D** and **E**).

**Figure 3.**
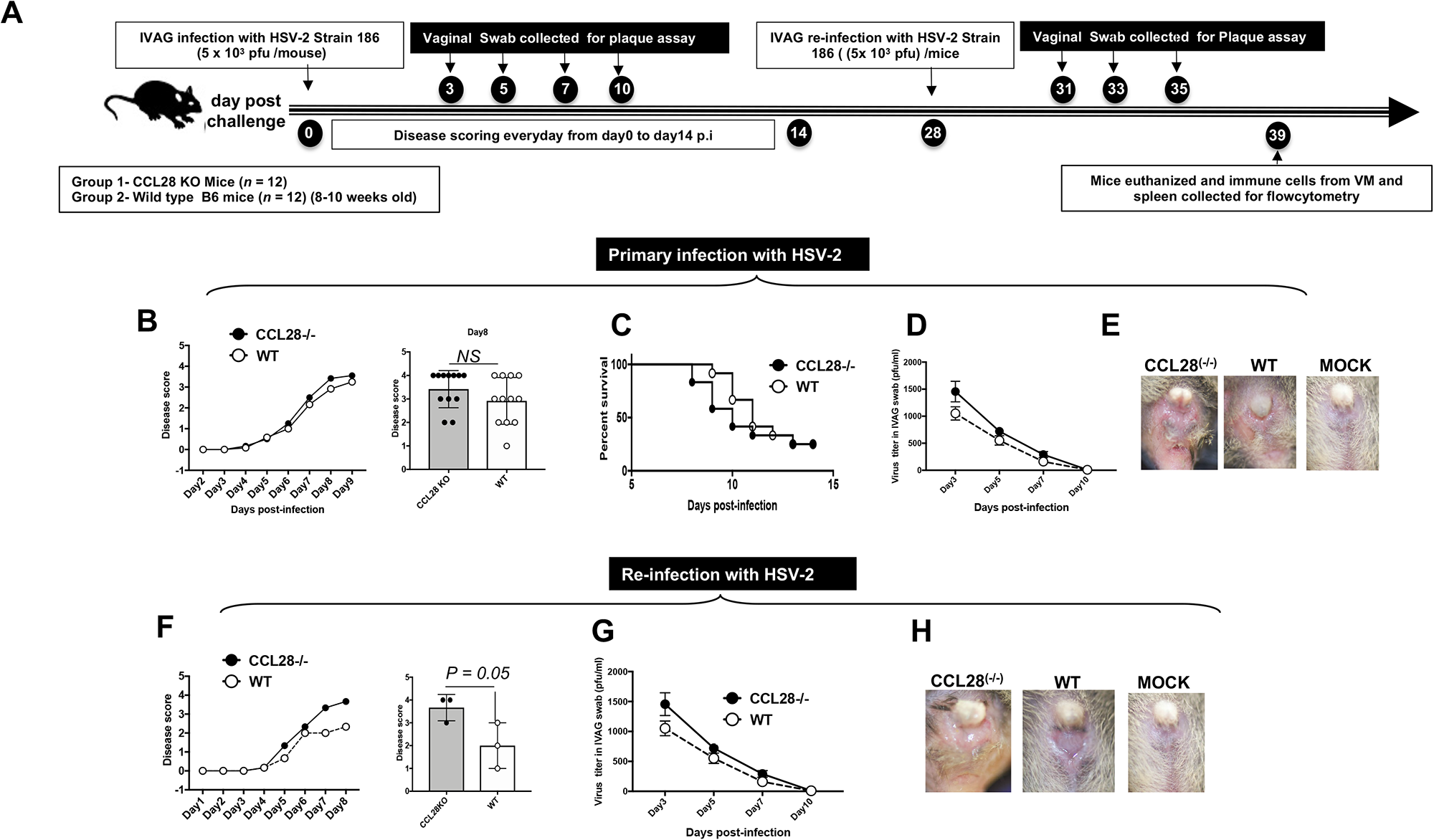
Susceptibility of CCL28^(-/-)^ knockout mice and B6 wild-type mice to genital herpes infection and disease following intravaginal infection and re-infection with HSV-2. (**A**) CCL28 KO mice (*n* = 12) and WT B6 mice (*n* = 12) were infected with IVAG with 5 x 10^3^ pfu of HSV-2 (strain 186). CCL28 KO and WT B6 mice were scored every day for 8 to 9 days p. I for symptoms of genital herpes and severity of genital herpes scored, as described in *Material and Methods.* The disease was scored as 0-no disease, 2-swelling and redness of external vagina, 3-severe swelling and redness of vagina and surrounding tissue and hair loss in the genital area, 4-ulceration and hair loss in the genital and surrounding tissue. The vaginal swabs were collected on days 3, 5, and 7 p. I to determine virus titers. (**B**) Disease scoring in CCL28 KO mice (*CCL28^-/-^*) (*n* = 12) and WT B6 mice (*WT*) (*n* = 12) was determined for 9 days after primary infection with HSV-2 strain 186 (*left panel).* The maximal disease severity in CCL28 KO mice (*CCL28^-/-^*) and WT B6 mice (*WT*) was determined 8 days after primary infection with HSV-2 strain 186(*right panel)*. (**C**) Survival graph of in CCL28 KO mice (*CCL28^-/-^*) and WT B6 mice (*WT*) determined for 14 days after primary infection with HSV-2. (**D**) The graph shows the virus titers detected in the vaginal swabs of CCL28 KO mice (*CCL28^-/-^*) and WT B6 mice (*WT*) collected on 3-, 5-, 7-, and 10-days post-primary infection with HSV-2. (**E**) Representative pictures of genital disease in CCL28 KO mice (*CCL28^-/-^*) and WT B6 mice (*WT*) taken on day 8 post-primary infection with HSV-2. (**F**) CCL28 KO mice (*n* = 3) and WT B6 mice (*n* = 3) were re-infected with IVAG with 5 x 10^3^ pfu of HSV-2 (strain 186) on day 28 post-primary infection. Disease scoring in CCL28 KO mice (*CCL28^-/-^*) and WT B6 mice (*WT*) was determined for 9 days after secondary re-infection with HSV-2 strain 186 (*left panel).* The maximal disease severity in CCL28 KO mice (*CCL28^-/-^*) and WT B6 mice (*WT*) was determined 8 days after secondary re-infection with HSV-2 (*right panel)*. (**G**) The graph shows the virus titers detected in the vaginal swabs of CCL28 KO mice (*CCL28^-/-^*) and WT B6 mice (*WT*) collected 5-, 7-, and 10-day post-secondary infection with HSV-2. (**H**) Representative pictures of genital disease in CCL28 KO mice (*CCL28^-/-^*) and WT B6 mice (*WT*) taken on day 8 post-secondary infection with HSV-2. The results are representative of two independent experiments. The indicated *P* values were calculated using the unpaired t-test and compared results obtained from CCL28 KO mice (*CCL28^-/-^*) and WT B6 mice (*WT*).

We further determined a potential role of CCL28 chemokine in genital herpes immunity following recall of memory immune responses. The CCL28 knockout mice (CCL28***^(-/-)^*** mice) and WT mice (*n* =3) were subject to a second IVAG infection with 5 x 10^3^ pfu of HSV-2 (strain 186 delivered on day 28 post-primary infection) (**Fig. 3A**). On day 28 post-primary infection, some animals (*n* = 3) were re-infected once. Mice were scored every day for 14 days p.i. for the severity of genital herpes, survival, and virus replication. Following the reinfection with HSV-2, we observed a significant increase in disease severity in CCL28***^(-/-)^*** mice compared to WT mice detected on day 8 post-re-infection (*P* = 0.05, **Fig. 3F** and **H**). Moreover, compared to WT mice, there was a significant increase in virus replication measured by plaque assay in vaginal swabs collected in the CCL28***^(-/-)^***mice at days 3, 5, 7, and 10 post-re-infection (*P* < 0.05, **Fig. 3G**).

These results: (*i*) Demonstrate a functional consequence of CCL28 deficiency that led to severe genital herpes disease caused by HSV-2 re-infection; (*ii*) Confirm that CCL28 mucosal chemokine plays an important role in protective immunity against genital herpes infection and disease.

### 4. CCL28 deficiency is associated with decreased frequencies of both CCR10^+^CD4^+^ and CCR10^+^CD8^+^ T cells within the vaginal mucosa following HSV-2 infection and re-infection

We next examined whether CCL28 deficiency, which was associated with severe genital herpes and increased virus replication following HSV-2 re-infection (**Fig. 3** above), would be the consequence of lower frequencies of CD4^+^ and CD8^+^ T cells within the VM. CCL28 knockout mice (CCL28^(-/-)^mice) and WT mice (n =12) were IVAG infected on day 0 with 5 x 10^3^ pfu of HSV-2 (strain 186) and then re-infected on day 28 with 5 x 10^3^ pfu HSV-2 strain186. On day 10 post-re-infection, mice were euthanized and cell suspensions from VM and spleen were analyzed by flow cytometry for the frequency of CCR10^+^CD4^+^ and CCR10^+^CD8^+^ T cells. As shown in **Fig. 4A**, we detected significantly lower frequencies of CD8^+^ T cells (P = 0.01, left panels) and CD4^+^ T cells (P < 0.01, right panels) in the VM of CCL28 knockout mice (CCL28^(-/-)^mice) compared to WT mice following re-infection with HSV-2. Moreover, we detected significantly lower frequencies of CCR10^+^ T cells (P = 0.007, top panels), CCR10^+^CD8^+^ T cells (P = 0.02, middle panels), and CCR10^+^CD4^+^ T cells (P = 0.02, bottom panels) in the VM of CCL28 knockout mice (CCL28^(-/-)^mice) compared to WT mice following re-infection with HSV-2 (**Fig. 4B**). The CCL28 deficiency specifically affected the frequencies of CCR10^+^ T cells, CCR10^+^CD8^+^ T cells and CCR10^+^CD4^+^ T cells within the VM (left panels) but not within the spleen (right panels).

**Figure 4.**
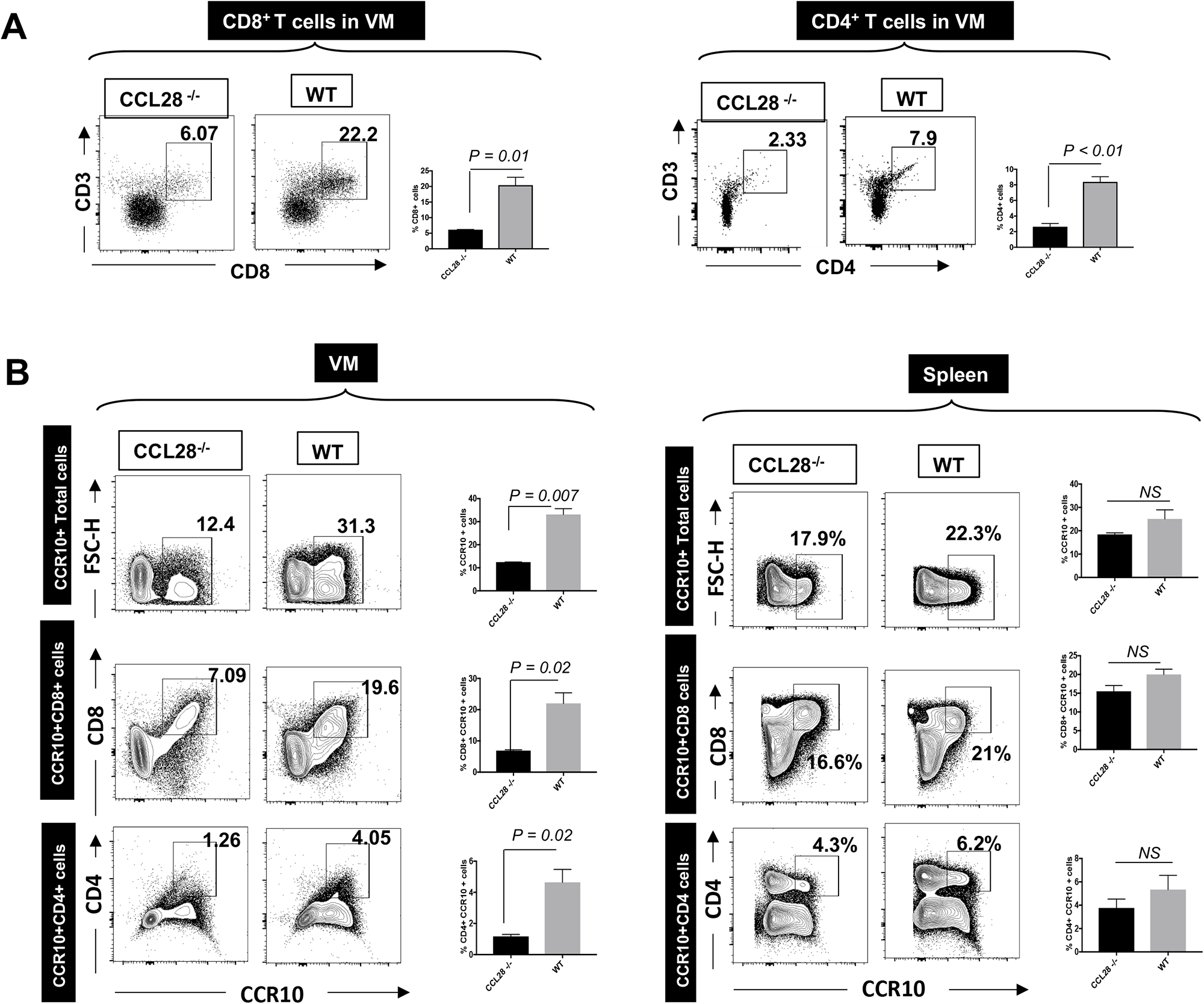
Frequencies of CD8^+^ and CD4^+^ T cells expressing CCR10, the receptor of CCL28, in the vaginal mucosa of CCL28^(-/-)^ knockout mice and B6 wild-type mice following intravaginal infection and re-infection with HSV-2. CCL28 KO mice (*CCL28^-/-^*) and WT B6 mice (*n* = 20) were IVAG infected with 5 x 10^3^ pfu of HSV-2 strain 186 and then re-infected with 5 x 10^3^ pfu of the same strain of HSV-2 on day 28 p.i. On day 10 post-final and secondary infection, mice were euthanized, and cell suspension from the vaginal mucosa (VM) and spleen was analyzed by flow cytometry for frequencies of CD8^+^ and CD4^+^ T cells expressing CCR10, the receptor of CCL28. (**A**) Representative and average frequencies of total CD8^+^ T cells (*left panels*) and total CD4^+^ T cells (*right panels*) in the VM of CCL28 KO mice (*CCL28^-/-^*) (*n* = 3) and WT B6 mice (*n* = 3) 10 days following re-infection with HSV-2. (**B**) Average frequencies of total CCR10^+^ T cells (*top panels*), CCR10^+^CD8^+^ T cells (*middle panels*), and CCR10^+^CD4^+^ T cells (*bottom panels*) detected in the VM (*right panels*) and spleen (*left panels*) of CCL28 KO mice (*CCL28^-/-^*) and WT B6 mice 10 days following re-infection with HSV-2. The results are representative of two independent experiments. The indicated *P* values were calculated using the unpaired *t*-test and compared results obtained from CCL28^(-/-)^ and WT mice.

These results: (*i*) demonstrate that CCL28 deficiency is associated with decreased frequencies of CCR10^+^CD4^+^ and CCR10^+^CD8^+^ T cells specifically in the vaginal mucosa (not in the spleen) following HSV-2 infection and re-infection; and (*ii*) suggest that CCL28 mucosal chemokine plays a critical role in the mobilization of protective memory CCR10^+^CD4^+^ and CCR10^+^CD8^+^ T cells, which express the CCR10 receptor of CCL28 chemokine, into the infected VM which likely protects locally against genital herpes infection and disease.

### 5. CCL28 deficiency is associated with decreased frequencies of effector memory CCR10^+^CD8^+^ T_EM_ cell subset, but not of central memory CCR10^+^CD8^+^ T_CM_ cell subset, within the vaginal mucosa following HSV-2 re-infection

We next examined whether CCL28 deficiency would affect the frequencies of specific subsets of memory CD4^+^ and CD8^+^ T cells within the VM, namely the effector memory T_EM_ and central memory T_CM_ cell subsets. CCL28 knockout mice (CCL28^(-/-)^ mice) and WT mice (n =12) were IVAG infected on day 0 with 5 x 10^3^ pfu of HSV-2 (strain 186) and then re-infected on day 28 with 5 x 10^3^ pfu HSV-2 strain186. On day 10 post-re-infection, mice were euthanized and cell suspensions from the VM and spleen were analyzed by flow cytometry for the frequency of effector memory T_EM_ and central memory T_CM_ cell subsets of both CD4^+^ T cells and CD8^+^ T cells. As shown in **Fig. 5A**, significantly lower frequencies of total memory CD8^+^ T cells (P = 0.01, left panels) were detected in the VM of CCL28 knockout mice (CCL28^(-/-)^ mice) compared to WT mice following re-infection with HSV-2. Moreover, the CCL28 deficiency was associated with decreased frequencies of effector memory CCR10^+^CD8^+^ T_EM_ cell subset, but not of central memory CCR10^+^CD8^+^ T_CM_ cell subset, within the vaginal mucosa following HSV-2 re-infection (**Fig. 5A**). However, deficiency in CCL28 neither affected the frequencies of effector memory CCR10^+^CD4^+^ T_EM_ cell subset nor of central memory CCR10^+^CD4^+^ T_CM_ cell subset within the vaginal mucosa following re-infection with HSV-2 (**Fig. 5B**).

**Figure 5.**
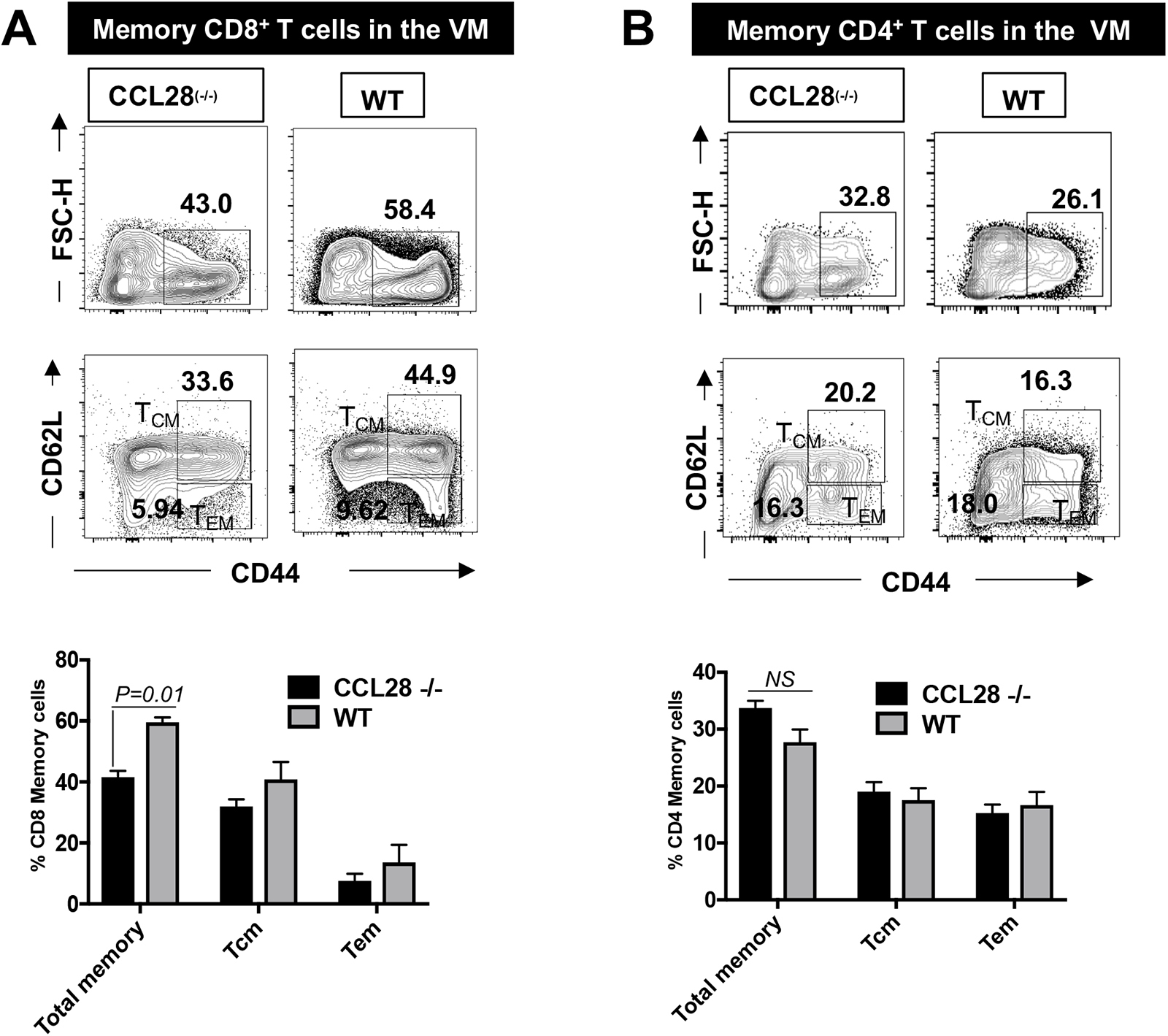
Frequencies of central and effector memory CD44^+^CD8^+^ and CD44^+^CD4^+^ T cells in the vaginal mucosa of CCL28^(-/-)^ knockout mice and B6 wild-type mice following intravaginal infection and re-infection with HSV-2. CCL28 KO mice (*CCL28^-/-^*) and WT B6 mice (*n* = 20) were IVAG infected with 5 x 10^3^ pfu of HSV-2 strain 186 and then re-infected with 5 x 10^3^ pfu of the same strain of HSV-2 on day 28 p.i. On day 10 post-re-infection, mice were euthanized and the frequencies of central memory CD44^+^CD62L^+^CD8^+^ T_CM_ cells and CD44^+^CD62L^+^CD4^+^ T_CM_ cells and of effector memory CD44^+^CD62L^-^CD8^+^ T_EM_ cells and CD44^+^CD62L^-^CD4^+^ T_EM_ cells were compared in the vaginal mucosa of CCL28 KO mice (*CCL28^-/-^*) and WT B6 mice using flow cytometry. (**A**) Representative data of the frequencies of total memory CD8^+^ T cells (*top 2 panels*) and central memory CD44^+^CD62L^+^CD8^+^ and effector memory CD44^+^CD62L^-^CD8^+^ T cells (*middle 2 panels*) in VM of CCL28 KO mice (*CCL28^-/-^*) and WT B6 mice re-infected with HSV-2. Average frequencies of total memory CD8^+^ T cells and central memory CD44^+^CD62L^+^CD8^+^T_CM_ cells and effector memory CD44^+^CD62L^-^CD8^+^ T_EM_ cells (*bottom panel*) in the VM of CCL28^-/-^ and WT B6 mice are re-infected with HSV-2. (**B**) Representative data of the frequencies of total memory CD4^+^ T cells (*top 2 panels*) and central memory CD44^+^CD62L^+^CD4^+^ and effector memory CD44^+^CD62L^-^CD4^+^ T cells (*middle 2 panels*) in VM of CCL28 KO mice (*CCL28^-/-^*) and WT B6 mice re-infected with HSV-2. Average frequencies of total memory CD4^+^ T cells and central memory CD44^+^CD62L^+^CD4^+^T_CM_ cells and effector memory CD44^+^CD62L^-^CD4^+^ T_EM_ cells (*bottom panel*) in the VM of CCL28^-/-^ and WT B6 mice are re-infected with HSV-2. The indicated *P* values were calculated using the unpaired *t*-test and compared results obtained from CCL28*^(-/-)^ (n = 3) and WT mice (n = 3) and the results are representative of two independent experiments*.

These results suggest that CCL28/CCR10 chemokine axis plays a major role in the mobilization of effector memory CCR10^+^CD44^+^ CD8^+^ T_EM_ cells within the VM site of herpes infection.

### 6. Decreased frequency of memory CD27^+^B220^+^ B cells in the vaginal mucosa of CCL28^(-/-)^ knockout mice compared to wild type B6 mice following HSV-2 infection and reinfection

Since antibodies and B cells also play a role in protection against genital herpes infection and disease, we finally examined whether CCL28 deficiency would affect the frequencies of total B cells and memory B cell subsets. CCL28 knockout mice (CCL28^(-/-)^ mice) and WT mice (n =12) were IVAG infected on day 0 with 5 x 10^3^ pfu of HSV-2 (strain 186) and then re-infected on day 28 with 5 x 10^3^ pfu HSV-2 strain186. On day 10 post-re-infection, mice were euthanized and cell suspensions from VM and spleen were analyzed by flow cytometry for the frequency of effector memory T_EM_ and central memory T_CM_ cell subsets of both CD4^+^ T cells and CD8^+^ T cells. There were significantly lower frequencies of CCR10^+^B220^+^ B cells (P = 0.01), CCR10^+^B220^+^CD27^+^ memory B cells (P = 0.05) were detected in the VM of CCL28^(-/-)^ mice compared to WT mice following re-infection with HSV-2 (**Fig. 6A**). As expected, the decrease in the frequencies of CCR10^+^B220^+^ B cells and CCR10^+^B220^+^CD27^+^ memory B cells specifically affected the CCR10 expressing B cells (P = 0.04, **Fig. 6B**). As shown in ELISPOT, the HSV-2-specific memory B cell response further confirmed a significant decrease in the function of HSV-specific memory B cells in CCL28^(-/-)^ mice compared to WT mice following re-infection with HSV-2 (P = 0.04, **Fig. 6C**). Our findings suggest that the CCL28/CCR10 chemokine axis functions through the infiltration of memory B cells to the site of re-activation, the VM.

**Figure 6.**
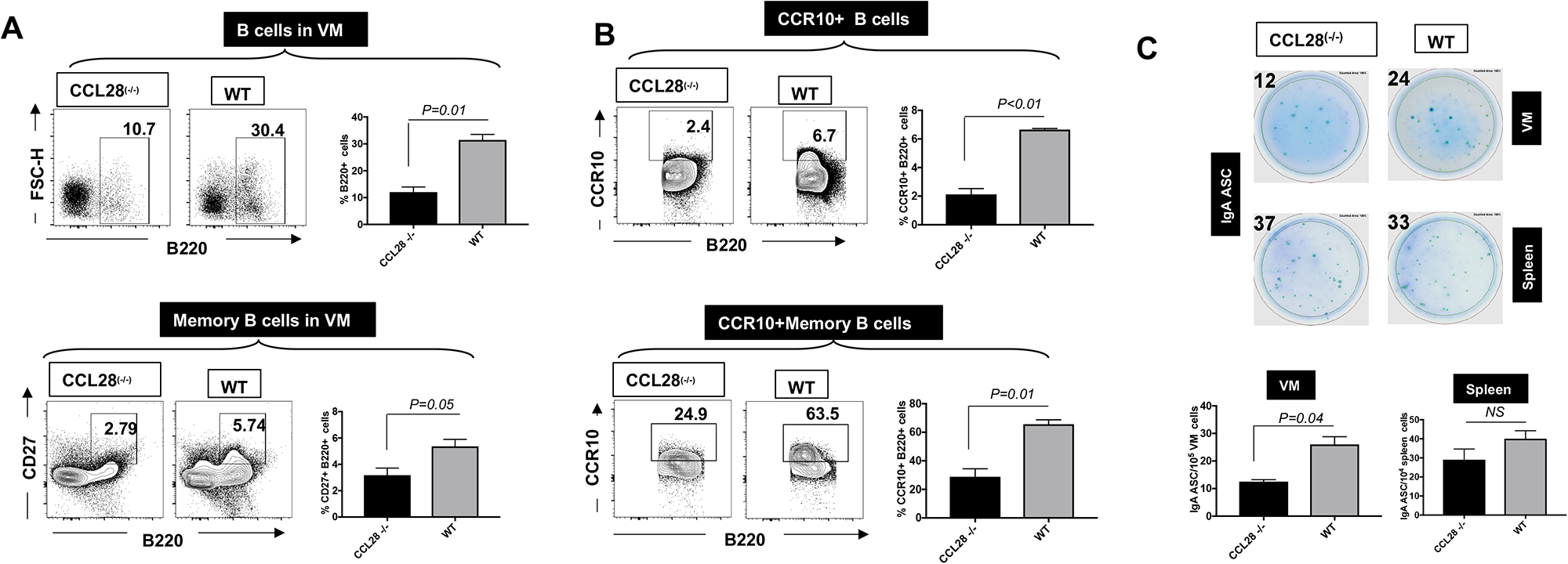
Frequencies of total B cells and memory B cells in the vaginal mucosa of CCL28^(-/-)^ knockout mice and B6 wild-type mice following intravaginal infection and re-infection with HSV-2. CCL28 KO mice (*CCL28^-/-^*) and WT B6 mice (*n* = 20) were IVAG infected with 5 x 10^3^ pfu of HSV-2 strain 186 and then re-infected with 5 x 10^3^ pfu of the same strain of HSV-2 on day 28 p.i. On day 10 post-re-infection, mice were euthanized and the frequencies of total B220^+^B cells and memory B220^+^B cells, expressing the expressing CCR10, the receptor of CCL28, were determined for flow cytometry in the VM and spleen of CCL28 KO mice and WT B6 mice. (**A**) Representative (*left 4 panels*) and average (*right 2 panels*) frequencies of total B220^+^B cells (*top 3 panels*) and memory B cells (*bottom 3 panels*) in VM of CCL28 KO mice (*n* = 3) and WT B6 mice (*n* = 3), 10 days following re-infection with HSV-2. (**B**) Representative (*left 4 panels*) and average (*right 2 panels*) frequencies of total B cells expressing CCR10, the receptor of CCL28, (CCR10^+^B220^+^ B cells *top 3 panels*), and of memory B cells expressing CCR10 (CCR10^+^B220^+^CD27^+^ memory B cells, *bottom 3 panels*) were determined in the VM of CCL28 KO mice and WT B6 mice 10 days following re-infection with HSV-2. (**C**) The ELISPOT images show IgA ASC in the VM (*top*) and spleen (*middle*) of CCL28 KO mice and WT B6 mice 10 days following re-infection with HSV-2. Corresponding average SFU for IgA ASC in the VM and Spleen are shown in the 2 *bottom panels*. The results are representative of two independent experiments. *P* values were calculated using the unpaired *t*-test and compared with results obtained in CCL28 KO mice and WT B6 mice.

These results suggest that: (*i*) CCL28/CCR10 chemokine axis affects the mobilization and function of memory CCR10^+^CD27^+^B220^+^ B cells, in addition to memory CCR10^+^CD44^+^ CD8^+^ T_EM_ cells, within the VM site of herpes infection; and (*ii*) CCL28 mucosal chemokine plays an important role in the mobilization of protective memory CCR10^+^CD8^+^ T_EM_ cells and CCR10^+^CD27^+^B220^+^ B cells, both expressing the CCR10 receptor of CCL28 chemokine, into the infected VM, which likely protect locally against genital herpes infection and disease.

## DISCUSSION

The four major mucosal-associated epithelial chemokines, CCL25, CCL28, CXCL14, and CXCL17, are expressed homeostatically in many mucosal tissues and play an important role in protecting mucosal surfaces from incoming infectious pathogens. Since the CCL28 mucosal chemokine is a chemoattractant for CCR10 expressing B and T cells and is highly expressed in the vaginal mucosa (VM), we investigated the role of the CCL28/CCR10 chemokine axis in the mobilization of HSV-specific memory B and T cells into VM site of herpes infection and its association with protection against genital herpes. We compared the differential expression of the CCR10, the receptor of CCL28, on herpes-specific CD8^+^ T cells from SYMP and ASYMP HSV-1 infected individuals using bulk RNA sequencing and flow cytometry. Genital herpes infection and disease were compared in CCL28 knockout (CCL28^(-/-)^) mice and wild-type B6 mice (WT) following genital herpes infection and re-infection with HSV-2 (strain186) genital infection. Frequencies of CCR10 expressing memory B and T cells within the VM were studied by flow cytometry and ELISPOT in SYMP and ASYMP HSV-1 infected mice. We found a significant increase in the frequencies of HSV-specific memory CD44^+^CD8^+^ T cells, expressing high levels of the CCR10 receptor, in herpes-infected ASYMP compared to SYMP individuals. Similarly, we detected significantly increased expression levels of the CCL28 chemokine in the VM of herpes-infected ASYMP mice compared to SYMP mice. Moreover, compared to WT mice, the CCL28 knockout (CCL28^(-/-)^) mice: (*i*) Appeared more susceptible to intravaginal infection and re-infection with HSV-2; (*ii*) Exhibited a decrease in frequencies of HSV-specific effector memory CCR10^+^CD44^+^ CD62L^-^ CD8^+^ T_EM_ cells, infiltrating the infected VM; and (*iii*) presented a decrease in the frequency of memory CD27^+^ B220^+^ B cells. Increased levels of CCL28 chemokine in asymptomatic herpes suggests a role of the CCL28/CCR10 mucosal chemokine axis in protection against genital herpes infection and disease through mobilization of high frequencies of both CCR10^+^B220^+^CD27^+^ memory B cells and HSV-specific memory CCR10^+^CD44^+^ memory CD8^+^ T cells within the infected vaginal mucosa.

Herpes simplex virus is one of the most common sexually transmitted viral infections worldwide (27). Globally, more women than men are infected by HSV-2 (28, 29), including ∼ 31 million in the U.S., and >300 million worldwide (30–32). Except for antiviral prophylaxis only available in developed countries, genital herpes simplex lacks effective treatment and there is no effective vaccination.

Studies that explore the correlates of protective immune response in HSV-infected but asymptomatic individuals would significantly aid in developing immune interventions to protect from herpes infection and disease in symptomatic patients. After the initial vaginal exposure, the virus replicates in vaginal epithelial cells (VEC), causing painful mucocutaneous blisters (33–39). Newly infected seronegative pregnant women can vertically transmit the virus to their newborns, causing encephalitis and death (40–42). Genital HSV-2 infection has also played a major role in driving the HIV prevalence (43–47), and there is no herpes vaccine or immunotherapy (27, 32, 48–50). Therefore, infected individuals rely on sustained or intermittent antiviral drugs (Acyclovir and derivatives), restrained sexual activity, and barrier methods to limit the spread of HSV-2 (51, 52).

In this study, we performed bulk RNA sequencing of herpes-specific CD8^+^ T cells isolated from PBMC of HSV-infected SYMP and ASYMP women. Our analysis revealed a unique differential regulation of the chemokine pathway and a significantly increased expression of CCR10 in ASYMP as compared to SYMP herpes-infected women. We further confirmed this result downstream by flow cytometry analysis of immune cells from PBMCs of SYMP and ASYMP HSV-infected women. Our results demonstrated an increased expression level of CCR10 on HSV-specific CD8^+^ T cells from ASYMP compared to SYMP women both transcriptionally and translationally. Based on these mRNA sequencing and flow cytometry results from SYMP and ASYMP HSV-infected women, we also explored the role of the mucosal chemokine CCL28/CCR10 chemokine axis in protection against genital herpes infection and disease using the mouse model. We used SYMP and ASYMP mice infected intravaginally with HSV and found an increased expression of CCL28 chemokine in the VM was associated with protection in ASYMP mice, but not in SYMP mice. We further confirmed this increased expression of chemokine CCL28 in VM of HSV-2 infected ASYMP mice by western blot and immunohistochemistry. The corresponding increase in the CCR10-expressing memory B and T cells was shown by flow cytometry, further suggesting a critical role of the mucosal chemokine CCL28/CCR10 chemokine axis in the protective immunity against genital herpes.

Chemokines are small, secreted polypeptides with chemotactic properties that regulate the trafficking of immune cells in homeostasis and inflammation (53). Inflammatory chemokines regulate inflammatory responses (54). Homeostatic chemokines are involved in T-cell immunity and immunopathology. They guide, attract, and relocate specific subsets of CD8+ T cells within and between lymphoid organs and non-lymphoid infected tissues (53, 55). Chemokines and their functions can be redundant and may not contribute to disease protection *in-vivo*. To further understand if CCL28 played a profound role associated with protection from disease severity in genital herpes, we used the CCL28^(-/-)^mice to understand if the absence of CCL28 can increase the severity of HSV-2 genital herpes. In addition, we studied whether the CCL28^(-/-)^ mice were more susceptible to genital re-infection with HSV-2 compared to WT mice, as reactivation of HSV-2 infection is the cause of recurrent genital herpes. *Since CCL28 chemokine appears to play a key role in the infiltration of memory immune cells into the VM compartment, we hypothesized that the recall of memory immune cells into the VM in re-infected CCL28 knock-out mice would be compromised.* It is also noteworthy to mention that the CCL28^(-/-)^ mice did not show any differences in disease or pathology during primary infection, but only showed increased susceptibility during re-infection. This could be due to CCL28 eliciting a better memory response by attracting more memory T cells to the site of infection during re-infection. This confirms previous reports showing that CCL28 regulates the migration of T cells that express the CCR10 receptor (56). CCL28 binds to both CCR10 and CXCR3, which are highly expressed on mucosal epithelia cells (56–65). The underlying mechanism of how the CCL28 improved the frequencies of antiviral CD8^+^ T cells in the VM is currently unknown. Nevertheless, our finding implies that delivering mucosal chemokines, such as CCL28, intravaginally using “mucosal tropic” adenovirus vectors in symptomatic mice could: (a) “re-open” this otherwise “immunologically closed compartment,” allowing infiltration by circulating CD8^+^ T cells; and/or (b) promote the formation, retention, and expansion of protective vaginal mucosa-resident CD8^+^ T_RM_ cells, which will suppress local HSV-2 replication, and hence prevent or reduce genital herpes disease. In future experiments, we will use AAV8 vectors expressing CCL28 mucosal chemokine that will be delivered intravaginally in HSV-2 infected mice, and examine recruitment, formation, retention, and expansion of HSV-specific CD8^+^ T_RM_ cells to the vaginal mucosa. We anticipate that sustained expression of CCL28 mucosal chemokine locally will be critical in mobilizing vaginal mucosal tissues-resident protective HSV-specific CD8^+^ T_RM_ cells that should control genital HSV-2 infection and disease. Those results will be the subject of a future report. Also, previous studies have shown that estrogen might have a crucial role in the protection against genital infection by regulating MEC/CCL28 expression in the uterus (58). The effect of sex hormones like estrogen on the functions of CCL28 will be an interesting area of research.

The immune profile of cells in the VM of infected mice showed that the CCL28^(-/-)^ mice had a decrease in CCR10 expressing CD8^+^ and CD4^+^ T cells and a decreased frequency of CCR10^+^CD44^+^ memory CD8^+^ T cells compared to WT mice. The role of the CCL28/CCR10 chemokine axis in the mobilization of IgA-secreting cells in mucosa has been well-established in the literature. To further understand if the CCL28 and its receptor have any role in humoral immunity during genital herpes infection, we studied the expression of CCR10 on B cells in VM. Interestingly, a majority of memory B cells in the VM of these mice expressed CCR10. There was also a decreased frequency of CD27^+^B220^+^ memory B cells in these CCL28^(-/-)^ mice. Increased frequency of CCR10 expressing CD8^+^ T cells in ASYMP herpes may suggest an association of mucosal chemokine CCL28 with protection in herpes infection. Thus, the mucosal chemokine CCL28 mediates protection from disease severity through the mobilization of both CCR10^+^CD44^+^ memory CD8^+^ T cells and CCR10^+^B220^+^CD27^+^ memory B cells to the VM. Recent studies have shown that low-dose CCL28 act as a molecular adjuvant when combined with the immunogen HSV-2 gB or HSV-2 gD with increased levels of virus-specific serum IgG and vaginal fluid IgA (66). This suggests that, in addition to the infiltration of memory T cells. CCL28 may also play a key role in the infiltration of memory B cells into the VM compartment.

During the last 20 years only a single vaccine strategy—adjuvanted recombinant HSV glycoprotein D (gD), with or without gB—has been tested and retested in clinical trials (67). Despite inducing strong HSV-specific neutralizing antibodies, this strategy failed to reach the primary endpoint of reducing herpes disease (68). These failures emphasize the need to induce T cell-mediated immunity (69). Following the resolution of viral infections, a long-lived memory CD8^+^ T cell subset that protects secondary (2°) infections is generated (18–22). This memory CD8^+^ T cell subset is heterogeneous but can be divided into three major subsets: (1) effector memory CD8^+^ T cells (CD8^+^ T_EM_ cells); (2) central memory CD8^+^ T cells (CD8^+^ T_CM_ cells); and (3) tissue-resident memory CD8^+^ T cells (CD8^+^ T_RM_ cells) (70). The three major sub-populations of memory T cells differ in their phenotype, function, and anatomic distribution. T_CM_ cells are CD62L^high^CCR7^high^CD103^low^. T_EM_ cells are CD62L^low^CCR7^low^CD103^low^. T_RM_ cells are CD62L^low^CCR7^low^CD103^high^CD11a^high^CD69^high^ (70–73). CD8^+^ T_RM_ cells are found in the vaginal mucosa and offer protection in mouse models of genital herpes (74). CD8^+^ T_EM_ cells are also found in the dermal-epidermal junction in women’s vaginal mucosa (75, 76). Once formed, T_RM_ cells do not re-enter the circulation and play an essential role in locally guarding mucosal tissues against secondary (2°) infections. However, the precise mechanisms by which non-circulating mucosa-resident memory CD8^+^ T_RM_ cells are formed, maintained, and expanded remain to be fully elucidated. In the present study, we found that a high frequency of CD8^+^ T_RM_ cells is retained in the vaginal mucosa of HSV-infected asymptomatic mice compared to symptomatic mice and that this is associated with CCL28 mucosal chemokine production. Specifically, we demonstrated that higher frequencies of vaginal mucosa tissue-resident antiviral memory CD8^+^ T cells (CD8^+^ T_RM_ cells) are a key mediator of protection against genital herpes, supporting previous reports (27, 75, 77–81). Since the primary cell target of HSV-2 is vaginal epithelial cells (VEC), the key to achieving anti-herpes mucosal immunity likely is to boost the frequencies of HSV-specific CD8^+^ T_RM_ cells in the vaginal mucosa that can expand locally and persist long-term. CD8^+^ T_RM_ cells persist long-term in tissues and are often embedded in the epithelial borders of mucosal tissues (82–86). However, little information exists on the mechanisms regulating the formation, retention, and expansion of vaginal-mucosa-resident CD8^+^ T_RM_ cells. To our knowledge, this report is the first to show CCL28/CCR10 chemokine axis mediated signals may be required for high frequencies of vaginal mucosa tissue-resident antiviral memory CD8^+^ T_RM_ cells. It remains to determine the mechanism of expansion and long-term retention of these CD8^+^ T_RM_ cells within the vaginal mucosa. Such knowledge will help design innovative vaccines to induce CD8^+^ T_RM_ cell-mediated protection from genital herpes. Collectively, this knowledge could greatly enhance our understanding of mucosal immunity and represents a unique opportunity to develop a powerful and long-lasting genital herpes vaccine that would have a significant impact on this disease’s epidemiology.

To our knowledge, our study represents the first in-depth analysis of the role of the CCL28/CCR10 chemokine axis in anti-herpes T and B cell responses in the VM during HSV-2 infection. We demonstrated that following intravaginal HSV-2 re-infection of B6 mice, high production of CCL28 chemokine in the VM was associated with increased infiltration of CCR10^+^CD44^+^ memory CD8^+^ T cells and CD27^+^B220^+^ memory B cells in the VM. Our findings could further aid in future innovative immunotherapeutic approaches for genital herpes.

## ACKNOWLEDGEMENTS

This work is supported by Public Health Service Research R01 Grants EY026103, EY019896, and EY024618 from the National Eye Institute (NEI), AI158060, AI150091, AI143348, AI147499, AI143326, AI138764, AI124911, and AI110902 from the National Institutes of Allergy and Infectious Diseases (NIAID) to LBM, and in part by The Discovery Center for Eye Research (DCER) and the Research to Prevent Blindness (RPB) grant. This work is dedicated to the memory of the late Professor Steven L. Wechsler “Steve” (1948–2016), whose numerous pioneering works on herpes infection and immunity laid the foundation for this line of research. We thank the NIH Tetramer Facility (Emory University, Atlanta, GA) for providing the Tetramers used in this study.

---

**<u>Supplementary Table 1</u>. Differentially expressed chemokine signaling pathway-specific genes in CD8^+^ T cell-specific to HSV-2 gB_561-569_ and VP11-12_220-228_ epitopes:** Shown are the log2 Fold Change and Adjusted P-values for the differentially expressed genes from Chemokine signaling pathway. The pairwise comparison of symptomatic and asymptomatic groups was performed using the DESeq2 package. Statistically, genes were considered differentially expressed when P < 0.5. and log2fold change >2.

## REFERENCES

1. Looker, K. J., A. S. Magaret, K. M. Turner, P. Vickerman, S. L. Gottlieb, and L. M. Newman. 2015. Global estimates of prevalent and incident herpes simplex virus type 2 infections in 2012. PLoS One 10: e114989.

2. Chentoufi, A. A., N. R. Dhanushkodi, R. Srivastava, S. Prakash, P. A. Coulon, L. Zayou, H. Vahed, H. A. Chentoufi, K. K. Hormi-Carver, and L. BenMohamed. 2022. Combinatorial Herpes Simplex Vaccine Strategies: From Bedside to Bench and Back. Front Immunol 13: 849515.

3. Harandi, A. M., B. Svennerholm, J. Holmgren, and K. Eriksson. 2001. Differential roles of B cells and IFN-gamma-secreting CD4(+) T cells in innate and adaptive immune control of genital herpes simplex virus type 2 infection in mice. J Gen Virol 82: 845–853.

4. MasCasullo, V., E. Fam, M. J. Keller, and B. C. Herold. 2005. Role of mucosal immunity in preventing genital herpes infection. Viral Immunol 18: 595–606.

5. Singh, R., A. Kumar, W. D. Creery, M. Ruben, A. Giulivi, and F. Diaz-Mitoma. 2003. Dysregulated expression of IFN-gamma and IL-10 and impaired IFN-gamma-mediated responses at different disease stages in patients with genital herpes simplex virus-2 infection. Clin Exp Immunol 133: 97–107.

6. Whitley, R. J., and R. L. Miller. 2001. Immunologic approach to herpes simplex virus. Viral Immunol 14: 111–118.

7. Inagaki-Ohara, K., T. Daikoku, F. Goshima, and Y. Nishiyama. 2000. Impaired induction of protective immunity by highly virulent herpes simplex virus type 2 in a murine model of genital herpes. Arch Virol 145: 1989–2002.

8. Celum, C. L. 2004. The interaction between herpes simplex virus and human immunodeficiency virus. Herpes 11 Suppl 1: 36A–45A.

9. Duerst, R. J., and L. A. Morrison. 2003. Innate immunity to herpes simplex virus type 2. Viral Immunol 16: 475–490.

10. Nagot, N., A. Ouedraogo, M. C. Defer, R. Vallo, P. Mayaud, and P. Van de Perre. 2007. Association between bacterial vaginosis and Herpes simplex virus type-2 infection: implications for HIV acquisition studies. Sex Transm Infect 83: 365–368.

11. Carr, D. J., and L. Tomanek. 2006. Herpes simplex virus and the chemokines that mediate the inflammation. Curr Top Microbiol Immunol 303: 47–65.

12. Mossman, K. L., P. F. Macgregor, J. J. Rozmus, A. B. Goryachev, A. M. Edwards, and J. R. Smiley. 2001. Herpes simplex virus triggers and then disarms a host antiviral response. J Virol 75: 750–758.

13. Koelle, D. M., and L. Corey. 2003. Recent progress in herpes simplex virus immunobiology and vaccine research. Clin Microbiol Rev 16: 96–113.

14. Parr, M. B., and E. L. Parr. 1998. Mucosal immunity to herpes simplex virus type 2 infection in the mouse vagina is impaired by in vivo depletion of T lymphocytes. J Virol 72: 2677–2685.

15. Kesson, A. M. 2001. Management of neonatal herpes simplex virus infection. Paediatr Drugs 3: 81–90.

16. Parra-Sanchez, M. 2019. Genital ulcers caused by herpes simplex virus. Enferm Infecc Microbiol Clin (Engl Ed) 37: 260–264.

17. Parr, M. B., L. Kepple, M. R. McDermott, M. D. Drew, J. J. Bozzola, and E. L. Parr. 1994. A mouse model for studies of mucosal immunity to vaginal infection by herpes simplex virus type 2. Lab Invest 70: 369–380.

18. Ariotti, S., M. A. Hogenbirk, F. E. Dijkgraaf, L. L. Visser, M. E. Hoekstra, J. Y. Song, H. Jacobs, J. B. Haanen, and T. N. Schumacher. 2014. T cell memory. Skin-resident memory CD8(+) T cells trigger a state of tissue-wide pathogen alert. Science 346: 101–105.

19. Mackay, L. K., A. Rahimpour, J. Z. Ma, N. Collins, A. T. Stock, M. L. Hafon, J. Vega-Ramos, P. Lauzurica, S. N. Mueller, T. Stefanovic, D. C. Tscharke, W. R. Heath, M. Inouye, F. R. Carbone, and T. Gebhardt. 2013. The developmental pathway for CD103CD8 tissue-resident memory T cells of skin. Nat Immunol.

20. Mackay, L. K., A. Rahimpour, J. Z. Ma, N. Collins, A. T. Stock, M. L. Hafon, J. Vega-Ramos, P. Lauzurica, S. N. Mueller, T. Stefanovic, D. C. Tscharke, W. R. Heath, M. Inouye, F. R. Carbone, and T. Gebhardt. 2013. The developmental pathway for CD103(+)CD8+ tissue-resident memory T cells of skin. Nature immunology 14: 1294–1301.

21. Wakim, L. M., A. Woodward-Davis, R. Liu, Y. Hu, J. Villadangos, G. Smyth, and M. J. Bevan. 2012. The molecular signature of tissue resident memory CD8 T cells isolated from the brain. J Immunol 189: 3462–3471.

22. Gebhardt, T., and L. K. Mackay. 2012. Local immunity by tissue-resident CD8(+) memory T cells. Front Immunol 3: 340.

23. Uematsu, S., and S. Akira. 2007. Toll-like receptors and Type I interferons. J Biol Chem 282: 15319–15323.

24. Gill, N., P. M. Deacon, B. Lichty, K. L. Mossman, and A. A. Ashkar. 2006. Induction of innate immunity against herpes simplex virus type 2 infection via local delivery of Toll-like receptor ligands correlates with beta interferon production. J Virol 80: 9943–9950.

25. Thapa, M., R. S. Welner, R. Pelayo, and D. J. Carr. 2008. CXCL9 and CXCL10 expression are critical for control of genital herpes simplex virus type 2 infection through mobilization of HSV-specific CTL and NK cells to the nervous system. J Immunol 180: 1098–1106.

26. Thapa, M., W. A. Kuziel, and D. J. Carr. 2007. Susceptibility of CCR5-deficient mice to genital herpes simplex virus type 2 is linked to NK cell mobilization. J Virol 81: 3704–3713.

27. Zhang, X., A. A. Chentoufi, G. Dasgupta, A. B. Nesburn, M. Wu, X. Zhu, D. Carpenter, S. L. Wechsler, S. You, and L. BenMohamed. 2009. A genital tract peptide epitope vaccine targeting TLR-2 efficiently induces local and systemic CD8+ T cells and protects against herpes simplex virus type 2 challenge. Mucosal Immunol 2: 129–143.

28. Haddow, L. J., E. A. Sullivan, J. Taylor, M. Abel, A. L. Cunningham, S. Tabrizi, and A. Mindel. 2007. Herpes simplex virus type 2 (HSV-2) infection in women attending an antenatal clinic in the South Pacific island nation of Vanuatu. Sexually transmitted diseases 34: 258–261.

29. Looker, K. J., G. P. Garnett, and G. P. Schmid. 2008. An estimate of the global prevalence and incidence of herpes simplex virus type 2 infection. Bull World Health Organ 86: 805–812, A.

30. Kalantari-Dehaghi, M., S. Chun, A. A. Chentoufi, J. Pablo, L. L., G. Dasgupta, D. M., A. Jasinskas, R. Nakajimi-sasaki, J. Felgner, G. Hermanson, L. BenMohamed, P. L. Felgner, and H. D. H. 2012. Discovery of potential diagnostic and vaccine antigens in herpes simplex virus-1 and -2 by proteome-wide antibody profiling. J. Virology In press.

31. Dervillez, X., S. Wechsler, A. B. Nesburn, and L. BenMohamed. 2012. Future of an “Asymptomatic” T-cell Epitope-Based Therapeutic Herpes Simplex Vaccine. Future Virology In Press.

32. Dasgupta, G., A. A. Chentoufi, M. Kalantari-Dehaghi, P. Falatoonzadeh, S. Chun, L. C. H., P. L. Felgner, H. D. H., L. BenMohamed. 2012. Immunodominant “Asymptomatic” Herpes Simplex Virus Type 1 and Type 2 Protein Antigens Identified by Probing Whole ORFome Microarrays By Serum Antibodies From Seropositive Asymptomatic Versus Symptomatic Individuals. J. Virology In press.

33. Peterslund, N. A. 1991. Herpesvirus infection: an overview of the clinical manifestations. Scand J Infect Dis Suppl 80: 15–20.

34. Rozenberg, F., C. Deback, and H. Agut. 2011. Herpes simplex encephalitis : from virus to therapy. Infect Disord Drug Targets 11: 235–250.

35. Nikkels, A. F., and G. E. Pierard. 2002. Treatment of mucocutaneous presentations of herpes simplex virus infections. Am J Clin Dermatol 3: 475–487.

36. Inoue, T., Y. Inoue, T. Nakamura, A. Yoshida, Y. Tano, Y. Shimomura, Y. Fujisawa, A. Aono, and K. Hayashi. 2002. The effect of immunization with herpes simplex virus glycoprotein D fused with interleukin-2 against murine herpetic keratitis. Jpn J Ophthalmol 46: 370–376.

37. Honda, M., and M. Niimura. 2000. [Alphaherpesvininae--dermatology]. Nippon rinsho 58: 895–900.

38. Liesegang, T. J. 1999. Classification of herpes simplex virus keratitis and anterior uveitis. Cornea 18: 127–143.

39. O’Brien, J. J., and D. M. Campoli-Richards. 1989. Acyclovir. An updated review of its antiviral activity, pharmacokinetic properties and therapeutic efficacy. Drugs 37: 233–309.

40. Whitley, R. J. 1994. Herpes simplex virus infections of women and their offspring: implications for a developed society. Proc Natl Acad Sci U S A 91: 2441–2447.

41. Westhoff, G. L., S. E. Little, and A. B. Caughey. 2011. Herpes simplex virus and pregnancy: a review of the management of antenatal and peripartum herpes infections. Obstet Gynecol Surv 66: 629–638.

42. Bettahi, I., X. Zhang, R. E. Afifi, and L. BenMohamed. 2006. Protective immunity to genital herpes simplex virus type 1 and type 2 provided by self-adjuvanting lipopeptides that drive dendritic cell maturation and elicit a polarized Th1 immune response. Viral Immunol 19: 220–236.

43. Mbopi-Keou, F. X., G. Gresenguet, P. Mayaud, H. A. Weiss, R. Gopal, M. Matta, J. L. Paul, D. W. Brown, R. J. Hayes, D. C. Mabey, and L. Belec. 2000. Interactions between herpes simplex virus type 2 and human immunodeficiency virus type 1 infection in African women: opportunities for intervention. The Journal of infectious diseases 182: 1090–1096.

44. Ouedraogo, A., N. Nagot, L. Vergne, I. Konate, H. A. Weiss, M. C. Defer, V. Foulongne, A. Sanon, J. B. Andonaba, M. Segondy, P. Mayaud, and P. Van de Perre. 2006. Impact of suppressive herpes therapy on genital HIV-1 RNA among women taking antiretroviral therapy: a randomized controlled trial. Aids 20: 2305–2313.

45. Freeman, E. E., H. A. Weiss, J. R. Glynn, P. L. Cross, J. A. Whitworth, and R. J. Hayes. 2006. Herpes simplex virus 2 infection increases HIV acquisition in men and women: systematic review and meta-analysis of longitudinal studies. Aids 20: 73–83.

46. Chentoufi, A. A., X. Dervillez, P. A. Rubbo, T. Kuo, X. Zhang, N. Nagot, E. Tuaillon, P. Van De Perre, A. B. Nesburn, and L. BenMohamed. 2012. Current trends in negative immuno-synergy between two sexually transmitted infectious viruses: HIV-1 and HSV-1/2. Current trends in immunology 13: 51–68.

47. Rubbo, P. A., E. Tuaillon, N. Nagot, A. A. Chentoufi, K. Bollore, J. Reynes, J. P. Vendrell, L. BenMohamed, and P. Van De Perre. 2011. HIV-1 infection impairs HSV-specific CD4(+) and CD8(+) T-cell response by reducing Th1 cytokines and CCR5 ligand secretion. Journal of acquired immune deficiency syndromes 58: 9–17.

48. Dasgupta, G., A. B. Nesburn, S. L. Wechsler, and L. BenMohamed. 2010. Developing an asymptomatic mucosal herpes vaccine: the present and the future. Future Microbiol 5: 1–4.

49. Zhang, X., F. A. Castelli, X. Zhu, M. Wu, B. Maillere, and L. BenMohamed. 2008. Gender-dependent HLA-DR-restricted epitopes identified from herpes simplex virus type 1 glycoprotein D. Clinical and vaccine immunology : CVI 15: 1436–1449.

50. Chentoufi, A. A., E. Kritzer, Y. M. D., A. B. Nesburn, and L. BenMohamed. 2012. Towards a Rational Design of an Asymptomatic Clinical Herpes Vaccine: The Old, The New, and The Unknown… Clinical and Developmental Immunology In Press.

51. Corey, L., A. Wald, R. Patel, S. L. Sacks, S. K. Tyring, T. Warren, J. M. Douglas, Jr., J. Paavonen, R. A. Morrow, K. R. Beutner, L. S. Stratchounsky, G. Mertz, O. N. Keene, H. A. Watson, D. Tait, and M. Vargas-Cortes. 2004. Once-daily valacyclovir to reduce the risk of transmission of genital herpes. The New England journal of medicine 350: 11–20.

52. Conant, M. A., D. W. Spicer, and C. D. Smith. 1984. Herpes simplex virus transmission: condom studies. Sexually transmitted diseases 11: 94–95.

53. Zlotnik, A., and O. Yoshie. 2012. The chemokine superfamily revisited. Immunity 36: 705–716.

54. Sallusto, F., and M. Baggiolini. 2008. Chemokines and leukocyte traffic. Nature immunology 9: 949–952.

55. Moser, B., M. Wolf, A. Walz, and P. Loetscher. 2004. Chemokines: multiple levels of leukocyte migration control. Trends Immunol 25: 75–84.

56. Wang, W., H. Soto, E. R. Oldham, M. E. Buchanan, B. Homey, D. Catron, N. Jenkins, N. G. Copeland, D. J. Gilbert, N. Nguyen, J. Abrams, D. Kershenovich, K. Smith, T. McClanahan, A. P. Vicari, and A. Zlotnik. 2000. Identification of a novel chemokine (CCL28), which binds CCR10 (GPR2). J Biol Chem 275: 22313–22323.

57. Castelletti, E., S. Lo Caputo, L. Kuhn, M. Borelli, J. Gajardo, M. Sinkala, D. Trabattoni, C. Kankasa, E. Lauri, A. Clivio, L. Piacentini, D. H. Bray, G. M. Aldrovandi, D. M. Thea, F. Veas, M. Nebuloni, F. Mazzotta, and M. Clerici. 2007. The mucosae-associated epithelial chemokine (MEC/CCL28) modulates immunity in HIV infection. PloS one 2: e969.

58. Cha, H. R., H. J. Ko, E. D. Kim, S. Y. Chang, S. U. Seo, N. Cuburu, S. Ryu, S. Kim, and M. N. Kweon. 2011. Mucosa-associated epithelial chemokine/CCL28 expression in the uterus attracts CCR10+ IgA plasma cells following mucosal vaccination via estrogen control. J Immunol 187: 3044–3052.

59. Eksteen, B., A. Miles, S. M. Curbishley, C. Tselepis, A. J. Grant, L. S. Walker, and D. H. Adams. 2006. Epithelial inflammation is associated with CCL28 production and the recruitment of regulatory T cells expressing CCR10. J Immunol 177: 593–603.

60. Lazarus, N. H., E. J. Kunkel, B. Johnston, E. Wilson, K. R. Youngman, and E. C. Butcher. 2003. A common mucosal chemokine (mucosae-associated epithelial chemokine/CCL28) selectively attracts IgA plasmablasts. Journal of immunology 170: 3799–3805.

61. O’Gorman, M. T., N. A. Jatoi, S. J. Lane, and B. P. Mahon. 2005. IL-1beta and TNF-alpha induce increased expression of CCL28 by airway epithelial cells via an NFkappaB-dependent pathway. Cellular immunology 238: 87–96.

62. Kunkel, E. J., C. H. Kim, N. H. Lazarus, M. A. Vierra, D. Soler, E. P. Bowman, and E. C. Butcher. 2003. CCR10 expression is a common feature of circulating and mucosal epithelial tissue IgA Ab-secreting cells. The Journal of clinical investigation 111: 1001–1010.

63. Meurens, F., M. Berri, J. Whale, T. Dybvig, S. Strom, D. Thompson, R. Brownlie, H. G. Townsend, H. Salmon, and V. Gerdts. 2006. Expression of TECK/CCL25 and MEC/CCL28 chemokines and their respective receptors CCR9 and CCR10 in porcine mucosal tissues. Veterinary immunology and immunopathology 113: 313–327.

64. Pan, J., E. J. Kunkel, U. Gosslar, N. Lazarus, P. Langdon, K. Broadwell, M. A. Vierra, M. C. Genovese, E. C. Butcher, and D. Soler. 2000. A novel chemokine ligand for CCR10 and CCR3 expressed by epithelial cells in mucosal tissues. Journal of immunology 165: 2943–2949.

65. Xiong, N., Y. Fu, S. Hu, M. Xia, and J. Yang. 2012. CCR10 and its ligands in regulation of epithelial immunity and diseases. Protein & cell 3: 571–580.

66. Yan, Y., K. Hu, M. Fu, X. Deng, S. Luo, L. Tong, X. Guan, S. He, C. Li, W. Jin, T. Du, Z. Zheng, M. Zhang, Y. Liu, and Q. Hu. 2021. CCL19 and CCL28 Assist Herpes Simplex Virus 2 Glycoprotein D To Induce Protective Systemic Immunity against Genital Viral Challenge. mSphere 6.

67. Kuo, T., C. Wang, T. Badakhshan, S. Chilukuri, and L. BenMohamed. 2014. The challenges and opportunities for the development of a T-cell epitope-based herpes simplex vaccine. Vaccine 32: 6733–6745.

68. Belshe, P. B., P. A. Leone, D. I. Bernstein, A. Wald, M. J. Levin, J. T. Stapleton, I. Gorfinkel, R. L. A. Morrow, M. G. Ewell, A. Stokes-Riner, G. Dubin, T. C. Heineman, J. M. Schulte, and C. D. Deal. 2012. Efficacy Results of a Trial of a Herpes Simplex Vaccine. The New England journal of medicine 366: 34–43.

69. Knipe, D. M., L. Corey, J. I. Cohen, and C. D. Deal. 2014. Summary and recommendations from a National Institute of Allergy and Infectious Diseases (NIAID) workshop on “Next Generation Herpes Simplex Virus Vaccines”. Vaccine 32: 1561–1562.

70. Gebhardt, T., P. G. Whitney, A. Zaid, L. K. Mackay, A. G. Brooks, W. R. Heath, F. R. Carbone, and S. N. Mueller. 2011. Different patterns of peripheral migration by memory CD4+ and CD8+ T cells. Nature 477: 216–219.

71. Mackay, L. K., L. Wakim, C. J. van Vliet, C. M. Jones, S. N. Mueller, O. Bannard, D. T. Fearon, W. R. Heath, and F. R. Carbone. 2012. Maintenance of T cell function in the face of chronic antigen stimulation and repeated reactivation for a latent virus infection. J Immunol 188: 2173–2178.

72. Mackay, L. K., L. Wakim, C. J. van Vliet, C. M. Jones, S. N. Mueller, O. Bannard, D. T. Fearon, W. R. Heath, and F. R. Carbone. 2012. Maintenance of T Cell Function in the Face of Chronic Antigen Stimulation and Repeated Reactivation for a Latent Virus Infection. J Immunol.

73. Mackay, L. K., A. T. Stock, J. Z. Ma, C. M. Jones, S. J. Kent, S. N. Mueller, W. R. Heath, F. R. Carbone, and T. Gebhardt. 2012. Long-lived epithelial immunity by tissue-resident memory T (TRM) cells in the absence of persisting local antigen presentation. Proc Natl Acad Sci U S A 109: 7037–7042.

74. Tang, V. A., and K. L. Rosenthal. 2010. Intravaginal infection with herpes simplex virus type-2 (HSV-2) generates a functional effector memory T cell population that persists in the murine genital tract. J Reprod Immunol 87: 39–44.

75. Zhu, J., T. Peng, C. Johnston, K. Phasouk, A. S. Kask, A. Klock, L. Jin, K. Diem, D. M. Koelle, A. Wald, H. Robins, and L. Corey. 2013. Immune surveillance by CD8alphaalpha+ skin-resident T cells in human herpes virus infection. Nature 497: 494–497.

76. Peng, T., J. Zhu, K. Phasouk, D. M. Koelle, A. Wald, and L. Corey. 2012. An effector phenotype of CD8+ T cells at the junction epithelium during clinical quiescence of herpes simplex virus 2 infection. J Virol 86: 10587–10596.

77. Zhang, X., X. Dervillez, A. A. Chentoufi, T. Badakhshan, I. Bettahi, and L. Benmohamed. 2012. Targeting the genital tract mucosa with a lipopeptide/recombinant adenovirus prime/boost vaccine induces potent and long-lasting CD8+ T cell immunity against herpes: importance of MyD88. J Immunol 189: 4496–4509.

78. Zhu, J., T. Peng, C. Johnston, K. Phasouk, A. S. Kask, A. Klock, L. Jin, K. Diem, D. M. Koelle, A. Wald, H. Robins, and L. Corey. 2013. Immune surveillance by CD8alphaalpha skin-resident T cells in human herpes virus infection. Nature.

79. Schiffer, J. T., L. Abu-Raddad, K. E. Mark, J. Zhu, S. Selke, D. M. Koelle, A. Wald, and L. Corey. 2010. Mucosal host immune response predicts the severity and duration of herpes simplex virus-2 genital tract shedding episodes. Proc Natl Acad Sci U S A 107: 18973–18978.

80. Zhu, J., F. Hladik, A. Woodward, A. Klock, T. Peng, C. Johnston, M. Remington, A. Magaret, D. M. Koelle, A. Wald, and L. Corey. 2009. Persistence of HIV-1 receptor-positive cells after HSV-2 reactivation is a potential mechanism for increased HIV-1 acquisition. Nat Med 15: 886–892.

81. Zhu, J., D. M. Koelle, J. Cao, J. Vazquez, M. L. Huang, F. Hladik, A. Wald, and L. Corey. 2007. Virus-specific CD8+ T cells accumulate near sensory nerve endings in genital skin during subclinical HSV-2 reactivation. J Exp Med 204: 595–603.

82. Laidlaw, B. J., N. Zhang, H. D. Marshall, M. M. Staron, T. Guan, Y. Hu, L. S. Cauley, J. Craft, and S. M. Kaech. 2014. CD4+ T cell help guides formation of CD103+ lung-resident memory CD8+ T cells during influenza viral infection. Immunity 41: 633–645.

83. Mackay, L. K., and T. Gebhardt. 2013. Tissue-resident memory T cells: local guards of the thymus. Eur J Immunol 43: 2259–2262.

84. Purwar, R., J. Campbell, G. Murphy, W. G. Richards, R. A. Clark, and T. S. Kupper. 2011. Resident memory T cells (T(RM)) are abundant in human lung: diversity, function, and antigen specificity. PloS one 6: e16245.

85. Sathaliyawala, T., M. Kubota, N. Yudanin, D. Turner, P. Camp, J. J. Thome, K. L. Bickham, H. Lerner, M. Goldstein, M. Sykes, T. Kato, and D. L. Farber. 2013. Distribution and compartmentalization of human circulating and tissue-resident memory T cell subsets. Immunity 38: 187–197.

86. Wu, T., Y. Hu, Y. T. Lee, K. R. Bouchard, A. Benechet, K. Khanna, and L. S. Cauley. 2014. Lung-resident memory CD8 T cells (TRM) are indispensable for optimal cross-protection against pulmonary virus infection. Journal of leukocyte biology 95: 215–224.

87. Rawls, W. E., D. Laurel, J. L. Melnick, J. M. Glicksman, and R. H. Kaufman. 1968. A search for viruses in smegma, premalignant and early malignant cervical tissues. The isolation of Herpesviruses with distinct antigenic properties. Am J Epidemiol 87: 647–655.

88. Segarra, T. J., E. Fakioglu, N. Cheshenko, S. S. Wilson, P. M. Mesquita, G. F. Doncel, and B. C. Herold. 2011. Bridging the gap between preclinical and clinical microbicide trials: blind evaluation of candidate gels in murine models of efficacy and safety. PLoS One 6: e27675.

89. Hendrickson, B. A., J. Guo, I. Brown, K. Dennis, D. Marcellino, J. Hetzel, and B. C. Herold. 2000. Decreased vaginal disease in J-chain-deficient mice following herpes simplex type 2 genital infection. Virology 271: 155–162.

