## Supplemental Figure 1 for "High Frequencies of Antiviral Effector Memory T_EM_ Cells and Memory B Cells Mobilized into Herpes Infected Vaginal Mucosa Associated With Protection Against Genital Herpes"

*Dhanushkodi et al Supplemental Fig. 1*

**A** Expression of CCL28 in the epithelia of human Vaginal mucosa

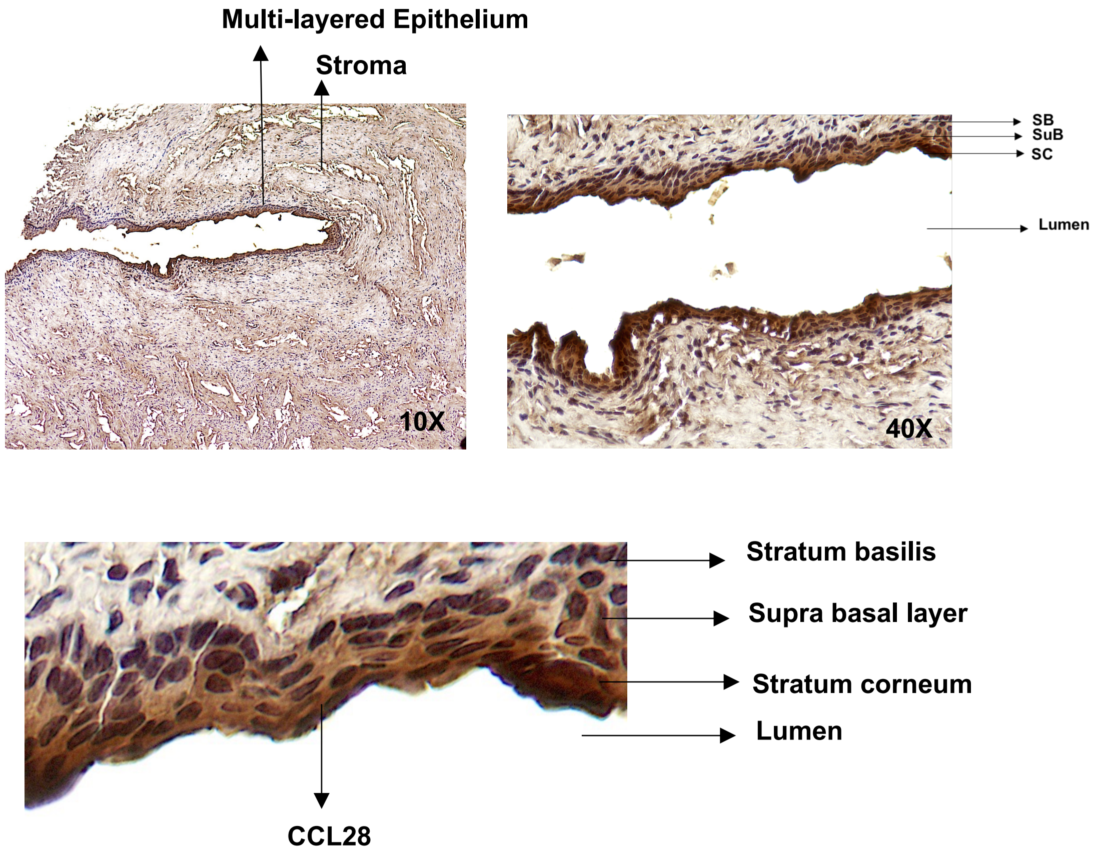
